# Amino acid substitutions associated with treatment failure of hepatitis C virus infection

**DOI:** 10.1101/2020.07.03.180240

**Authors:** María Eugenia Soria, Carlos García-Crespo, Brenda Martínez-Gónzalez, Lucía Vázquez-Sirvent, Rebeca Lobo-Vega, Ana Isabel de Ávila, Isabel Gallego, Qian Chen, Damir García-Cehic, Meritxell Llorens-Revull, Carlos Briones, Jordi Gómez, Cristina Ferrer-Orta, Nuria Verdaguer, Josep Gregori, Francisco Rodríguez-Frías, María Buti, Juan Ignacio Esteban, Esteban Domingo, Josep Quer, Celia Perales

**Affiliations:** Department of Clinical Microbiology, IIS-Fundación Jiménez Díaz, UAM. Av. Reyes Católicos 2, 28040 Madrid, Spain; Liver Unit, Internal Medicine Hospital Universitari Vall d’Hebron, Vall d’Hebron Institut de Recerca (VHIR), 08035, Barcelona, Spain; Centro de Biología Molecular “Severo Ochoa” (CSIC-UAM), Consejo Superior de Investigaciones Científicas (CSIC), Campus de Cantoblanco, 28049, Madrid, Spain; Centro de Investigación Biomédica en Red de Enfermedades Hepáticas y Digestivas (CIBERehd) del Instituto de Salud Carlos III, 28029, Madrid, Spain; Centro de Astrobiología (CAB, CSIC-INTA), 28850 Torrejón de Ardoz, Madrid, Spain; Instituto de Parasitología y Biomedicina ‘López-Neyra’ (CSIC), Parque Tecnológico Ciencias de la Salud, Armilla, 18016, Granada, Spain; Structural Biology Department, Institut de Biología Molecular de Barcelona CSIC, Barcelona, Spain; Roche Diagnostics, S.L., Sant Cugat del Vallés, 08174, Barcelona, Spain; Biochemistry and Microbiology Departments, VHIR-HUVH, Barcelona, Spain

**Keywords:** next-generation sequencing, viral quasispecies, viral fitness, antiviral agents, viral diagnostics, treatment planning

## Abstract

Despite the high virological response rates achieved with current directly-acting antiviral agents (DAAs) against hepatitis C virus (HCV), around 2% to 5% of treated patients do not achieve a sustained viral response. Identification of amino acid substitutions associated with treatment failure requires analytical designs, such as subtype-specific ultra-deep sequencing (UDS) methods for HCV characterization and patient management. Using this procedure, we have identified six highly represented amino acid substitutions (HRSs) in NS5A and NS5B of HCV from 220 patients who failed therapy, which are not *bona fide* resistance-associated substitutions (RAS). They were present frequently in basal and post-treatment virus of patients who failed therapy to different DAA-based therapies. Contrary to several RAS, HRSs belong to the acceptable subset of substitutions according to the PAM250 replacement matrix. Coherently, their mutant frequency, measured by the number of deep sequencing reads within the HCV quasispecies that encode the relevant substitutions, ranged between 90% and 100% in most cases. Also, they have limited predicted disruptive effects on the three-dimensional structures of the proteins harboring them. Possible mechanisms of HRS origin and dominance, as well as their potential predictive value of treatment response are discussed.

## Introduction

Hepatitis C virus (HCV) currently infects chronically around 71 million people worldwide (https://www.who.int/news-room/fact-sheets/detail/hepatitis-c), and it replicates as mutant clouds called viral quasispecies that confer an enormous adaptive potential to the virus [1–4]. Despite available direct-acting antiviral (DAA)-based therapies being extremely effective, in 2% to 5% of treated patients, viral load is not efficiently suppressed. Given the massive number of patients undergoing treatment worldwide, characterization of resistance-associated substitutions (RAS) has become part of HCV therapy management [5–8]. Selection of RAS associated with treatment failure is increasingly reported, concomitantly with the number of treated patients [9–13]. RAS can be also present in naïve patients who have not received DAA therapies, again as documented with several cohorts worldwide [14–18].

In a recent deep sequencing analysis of 220 subtyped HCV samples from infected patients who failed therapy, collected from 39 Spanish hospitals, we determined amino acid sequences of the DAA-target proteins NS3, NS5A and NS5B, by ultra-deep sequencing (UDS) of HCV patient samples, in search of RAS [9]. Interestingly, 18.6% of the patients that failed therapy did not include any substitutions that could be considered *bona fide* RAS, according to current guidelines [6]. Similar observations have been made with other patient cohorts [10,11,19–26]. This finding has raised the possibility that mechanisms other than RAS selection may contribute to treatment failure of HCV-infected patients.

Previous model studies with HCV replicating in human hepatoma Huh-7.5 cells indicated that such alternative mechanisms may exist. Specifically, studies with isogenic HCV populations (derived from the same initial genome) showed that the virus endowed with up to 2.3-fold increase in replicative fitness displayed increased resistance to several classes of anti-HCV inhibitors, including DAAs [27–29].

In the present study with HCV of chronically infected patients who failed DAA therapies, we document that a number of amino acid substitutions in NS5A and NS5B that are not *bona fide* RAS are present in basal samples (prior to DAA treatment), and remain dominant in the HCV quasispecies of a considerable proportion of patients. They have been termed highly represented substitutions (HRS), and can be found in isolation or in combination in the same viral sample. Their frequency is influenced by the viral genotype, and contrary to RAS, they belong to the statistically accepted class of substitutions in protein evolution, and predict minimal distortions in the structure of the corresponding proteins. The recognition of HRS provides new insights into the population dynamics of HCV *in vivo*, suggests that mechanisms other than RAS selection may contribute to treatment breakdown, and opens the possibility that HRS may be used as prognostic markers of DAA treatment response.

## Results

### Amino acid variations in HCV-infected patients failing DAA-based therapies. Defining highly represented substitutions (HRSs) in therapy outcome

RAS identified by UDS have been recently described in a cohort of 220 HCV-infected patients failing DAA therapies [9]. To provide a broad picture of all substitutions identified in NS3 (within amino acids 32 to 179), NS5A (within amino acids 24 to 152) and NS5B (within amino acids 124 to 320) proteins in these patients (Table S1), we constructed a heat map representing the frequency of each substitution in each viral sample (Figure 1A). We defined as highly represented those substitutions present in more than 20% of the patients. Contrary to the expectations, only three substitutions (Y93H in NS5A, as well as L159F and C316N in NS5B) out of the nine that fulfilled the frequency criterion belonged to the previously defined RAS [6,30–32] (Figure 1B). The three RAS were not considered part of the HRS for further analyses and calculations in the present study. The percentage of infected patients whose HCV carried any one of the HRS was statistically significant relative to those showing any other amino acid substitution within the protein regions analyzed. According to the proportion test, the range of p values was as follows: for T64A, p = 3.29×10^−14^ to 0.04; for R78K, p = 1.04×10^−11^ to 0.03; for S213C, p < 2.20×10^−16^ to 0.001; for A218S, p < 2.20×10^−16^ to 0.002; for S231N, p = 4.57×10^−15^ to 0.03; for Q309R, p = 5.57×10^−12^ to 0.04. Thus, HRSs outstand over other substitutions by their frequency among patients who failed DAA therapies.

**Figure 1.**
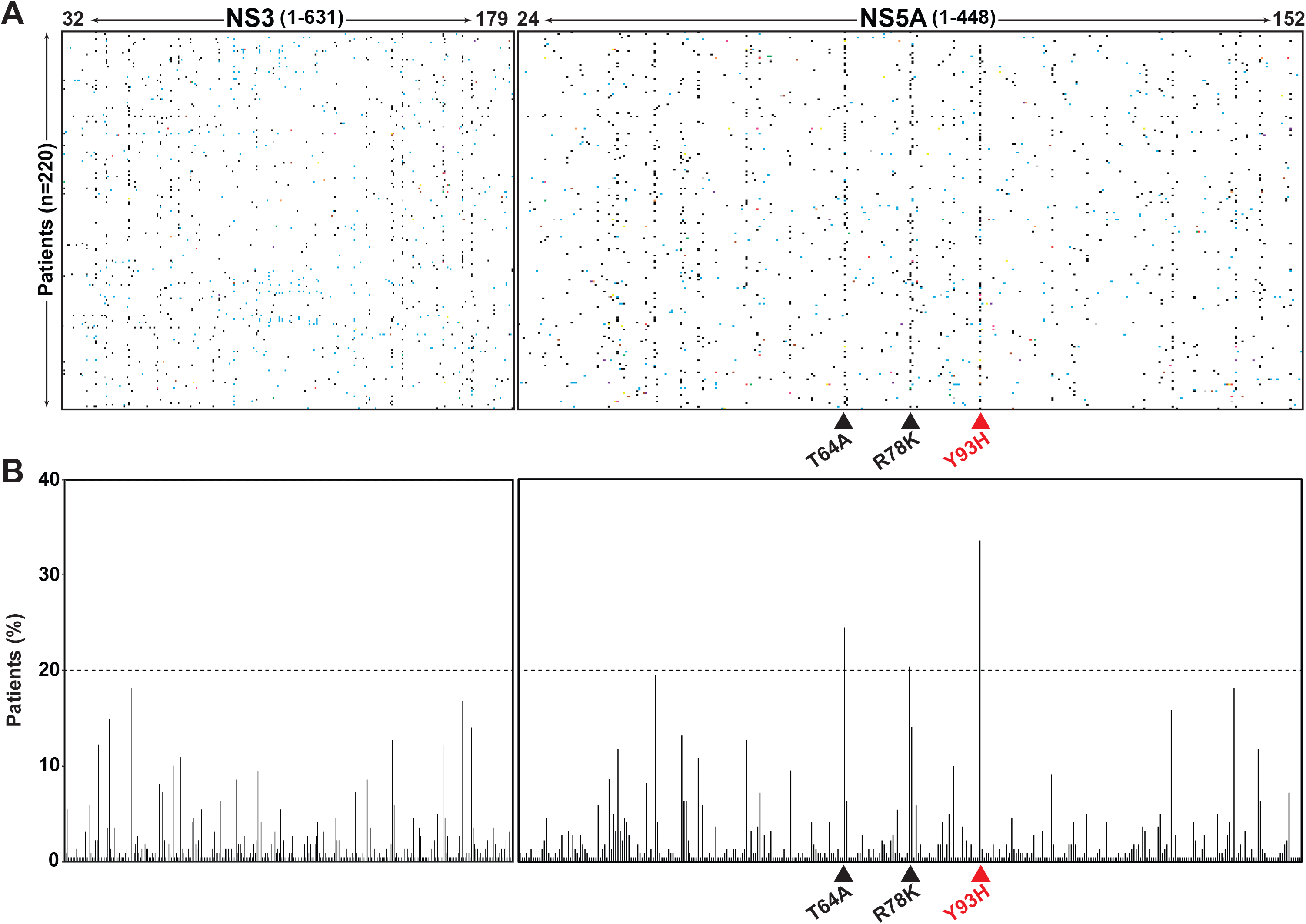
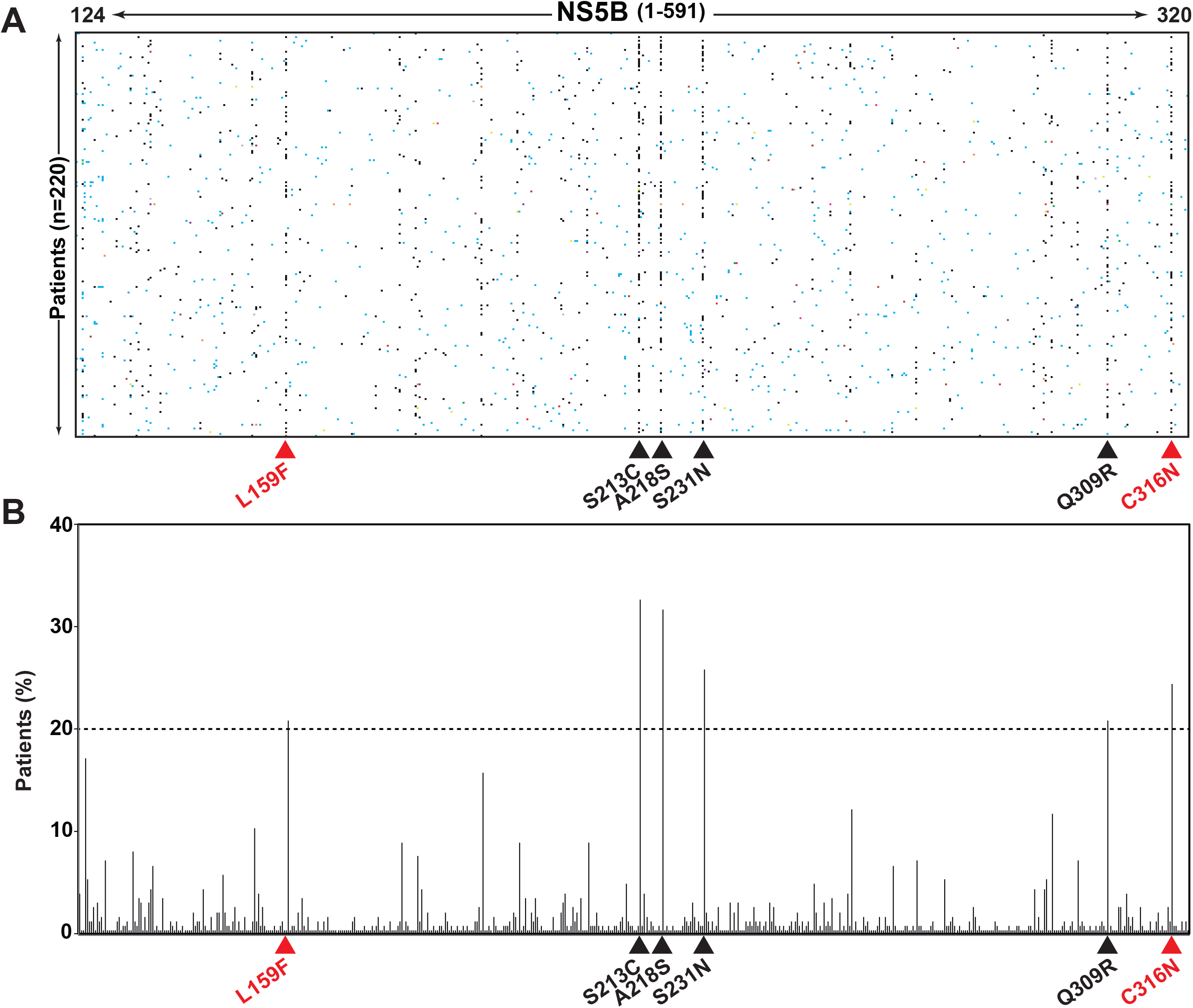
Heat map of amino acid substitutions and their distribution among patients, following treatment failure with DAAs. **(A)** The amino acid residues that were sequenced within proteins NS3, NS5A, and NS5B are indicated at the top. In parenthesis the protein length of NS3, NS5A and NS5B is indicated. Each horizontal dot alignment represents one of the total 220 patients analyzed (ordinate). Each vertical dot alignment corresponds to an amino acid position where an amino acid substitution was found; the substitution frequency in a sample (given by the proportion of reads with the relevant amino acid substitution) is denoted by the dot color: black (90.1-100%), grey (80.1-90%), pink (70.1-80%), purple (60.1-70%), red (50.1-60%), green (40.1-50%), orange (30.1-40%), yellow (20.1-30%), brown (10.1-20%), blue (1-10%) and white (<1%, below the limit of detection). The excel file including all amino acid frequencies represented by colors is available upon request. **(B)** Distribution of the amino acid substitutions depicted in A, according to the percentage of patients in whom each substitution was found (ordinate). The discontinuous horizontal line marks the 20% cut-off patient frequency used to define prevalent substitutions. The highly represented substitutions (HRSs) are indicated with a black triangle, and *bona fide* resistance-associated substitutions (RAS) with a red triangle, below the boxes. The complete list, location, statistical acceptability, and frequency among patients of all amino acid substitutions is given in Table S1.

### HRS dependence on viral subtype

Since HCV subtype can influence the response to treatment and RAS selection [9,10], it was interesting to explore if HCV subtype may also affect the types of HRSs found in the viral proteins. In NS5A, while T64A was present in patients infected with HCV of the three main viral genotypes (G1b, G1a and G3a), R78K was found mainly in G1a HCV-infected patients (Figure 2A). For G1a, S213C and A218S were not represented, and the order of the other HRSs according to the percentage of patients harboring them was R78K>Q309R>T64A>S231N. In contrast, for G1b, R78K was not present, and the order of abundance of other HRSs was S213C=A218S>S231N>T64A>Q309R. These differences of distribution among the two genotypes are statistically significant (for T64A, p=0.03899 and p=5.89×10^−4^ for the comparison between G1b and G1a or G3a, respectively; for R78K, S213C and A218S, p=2.2×10^−16^ for the comparisons between G1a and G1b; for S231N and Q309R, p=3.4×10^−14^ and p=5.9×10^−6^ for the comparisons between G1a and G1b, respectively; proportion test). These results suggest that the HRSs display a certain degree of subtype specificity, as has been previously reported for RAS [6,9,10].

**Figure 2.**
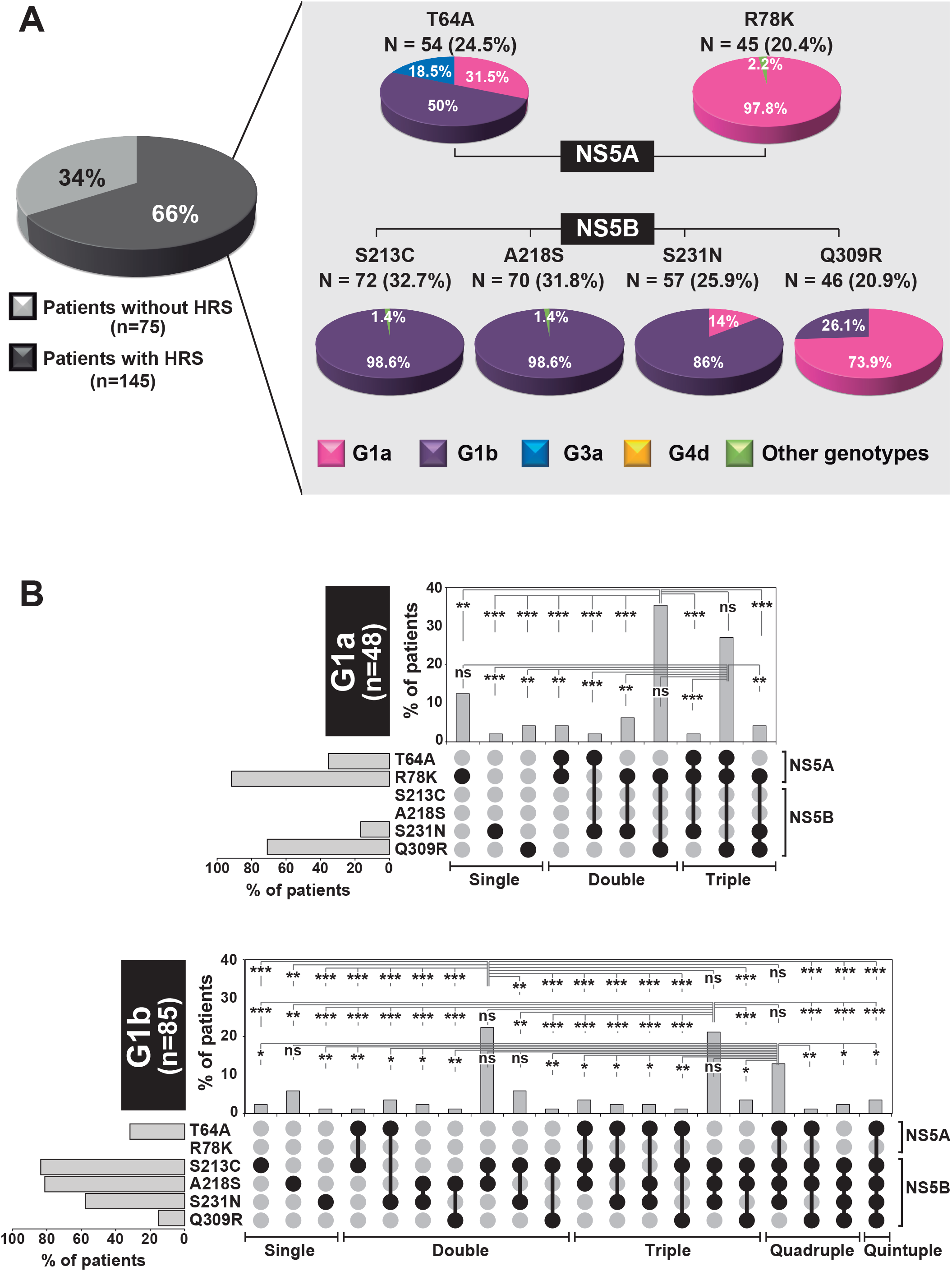
Distribution of single and combined HRSs among subtypes, following treatment failure with DAAs. **(A)** The circle on the left indicates the percentage of patients carrying at least one HRS. On the right the HRS distribution according to viral protein and HCV subtype is represented. The number in parenthesis indicates the percentage of patients carrying each HRS considering the 220 patient cohort. **(B)** Distribution of single and combined HRSs in the failure (post-treatment) samples. The display is divided into data for HCV genotype G1a (top bloc) and G1b (bottom bloc). For each bloc, the top panel indicates the percentage of patients (ordinate) in whom single or combined HRSs is found (individual or linked black circles below the panel). At the left of the HRS list, the grey horizontal bars depict the percentage of patients of each subtype (G1a and G1b) in whom each HRS is found; absence of a bar means that the HRS was absent. The protein where each HRS maps is shown on the right. It should be noted that the HRS associations are based on their presence in the same HCV sample, deduced from the same or different amplicons. Linkage of two HRSs in the same genomic molecule can be only indirectly inferred from their abundance in their corresponding amplicon population. Cases in which differences are statistically significant are indicated: ns=not significant; * p<0.05; ** p<0.01; *** p<0.001; proportion test. P values are indicated in the supplementary Table S2.

### Frequency of individual and combined HRSs in HCV quasispecies and among HCV-infected patients

The frequency of HRSs within the HCV mutant spectra of individual HCV samples was calculated from the percentage of UDS reads carrying each relevant mutation, taking into consideration the 1% limit of detection of amino acid substitutions [33]. Ninety-four percent of HRSs were found at frequencies that ranged between 90% and 100% in the viral quasispecies, whereas only 6% of them were found at frequencies between 1% and 89.9% (p < 2.2 × 10^−16^; proportion test) (Figure S1). Therefore, a high representation among patients paralleled a high frequency in the HCV mutant spectrum of each patient.

To study whether HRSs at NS5A and NS5B occur independently or they are preferentially combined in individual infected patients, the percentage of patients carrying HCV genomes with single, double, triple, quadruple and quintuple combinations was compared with the frequency expected from their individual frequency among patients (Figure 2B and Table S2). Application of Bayes theorem indicated that for genotype G1a there were no significant differences between observed and expected HRS associations (p=0.0589; chi-square test with Monte Carlo correction). For the most represented HRS combinations, the p values calculated with the proportion test were p=0.1732 for the R78K+Q309R, and p=0.2089 for T64A+R78K+Q309R. For HCV G1b no difference was evidenced by application of Bayes theorem (p=1.00; chi-square test with Monte Carlo correction). For the most represented combinations, the proportion test yielded p=0.2334 for the combination S213C+A218S, p=0.0551 for S213C+A218S+S231N and p=0.1047 for T64A+S213C+A218S+S231N. Thus, in the majority of cases the frequency of associated HRS is that expected from their individual o combined frequencies in infected patients.

### Prevalence of HRS in basal (prior to treatment) samples

To further investigate the possible origin of the HRSs found after treatment failure, we analyzed whether HRSs were already present before treatment implementation. To this aim, we had available 69 paired basal-failure samples that were analyzed following the same UDS procedure [9,33]. Seventy-four percent of the paired basal-failure samples carried at least one of the HRSs in the basal and/or post-treatment sample. For HCV G1a and G1b separately, a distinction was made for each HRS to indicate if its frequency within the viral quasispecies was equal or different between the basal and the post-treatment sample (Figure 3A and Table S3). The frequency of most HRSs was similar in the two paired samples. The results suggest that at least part of the HRSs found following treatment failure are dictated by their presence prior to treatment. Regarding the percentage of patients, the conclusion is also valid for HRS combinations found in basal and post-treatment samples (Figure 3B). A large proportion (62.5%) of double or higher order HRS combinations were present both in the basal and post-treatment HCV sample. Thus, the difference in HRSs and their combinations between the basal and the post-treatment sample was not statistically significant for patients infected with HCV G1a (p=0.5417; chi-square test with Monte Carlo correction), and virtually inexistent for those infected with HCV G1b (p=0.9925; chi-square test with Monte Carlo correction).

**Figure 3.**
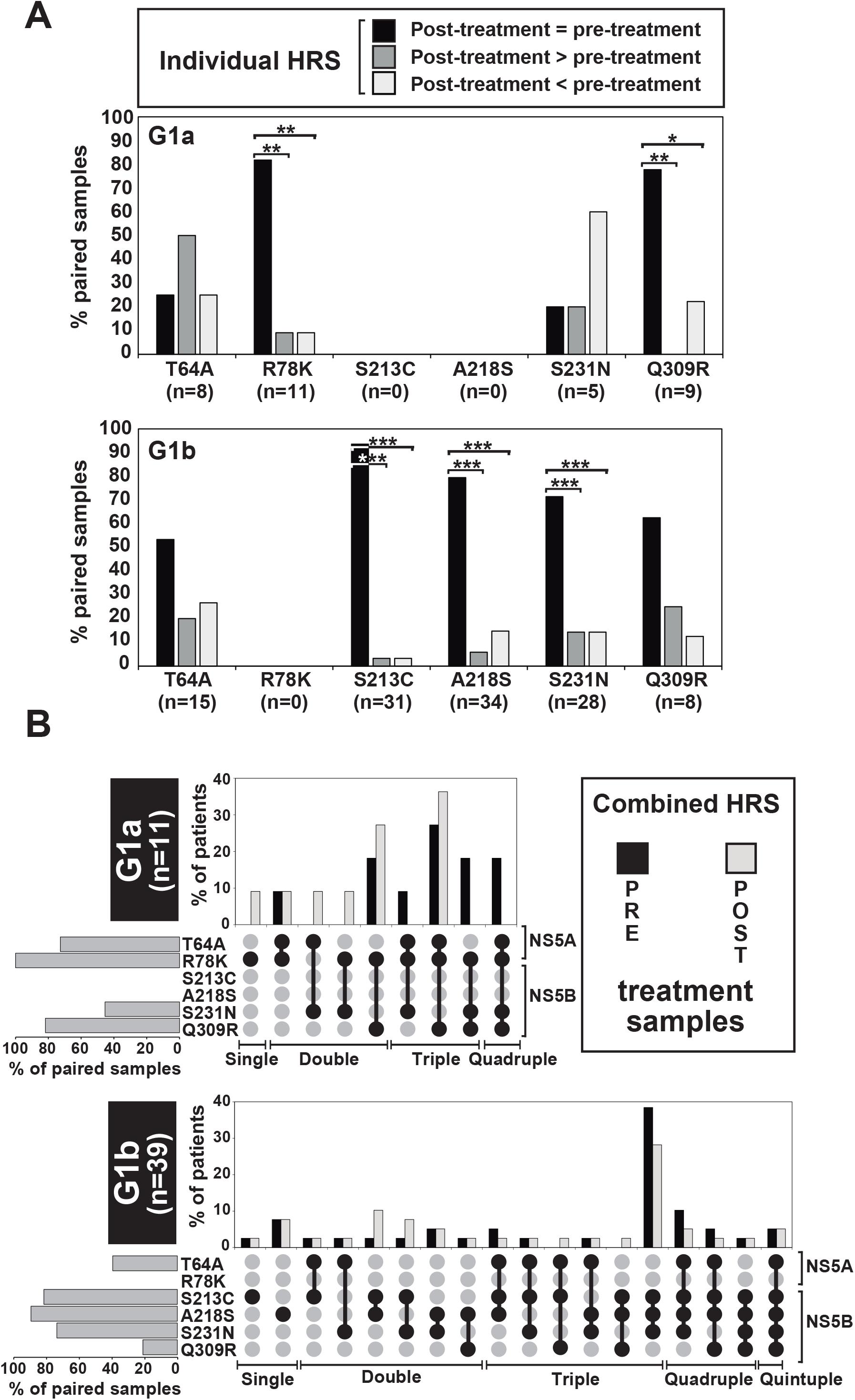
Comparison of the frequency of individual and combined HRSs in HCV from patients in the basal (pre-treatment) and post-treatment (failure) samples. **(A)** Distribution of the individual HRSs (given in abscissa) among patients (ordinate), according to their frequency in the post-treatment sample being equal (in all cases between 90%-100% within the viral quasispecies), higher or lower than in the pre-treatment sample (vertical bars with code in upper box). Cases in which differences are statistically significant are indicated: * p<0.05; ** p<0.01; *** p<0.001; proportion test. P values are indicated in the supplementary Table S3. **(B)** Distribution of single and combined HRSs in the basal (pre-treatment) and failure (post-treatment) samples (code in box on the right). The display is divided into data for HCV genotype G1a (top bloc) and G1b (bottom bloc). For each bloc, the top panel indicates the percentage of patients (ordinate) in whom single or combined HRSs is found (individual or linked black circles below the panel). At the left of the HRS list, the grey horizontal bars depict the percentage of patients of each subtype (G1a and G1b) in whom each HRS is found; absence of a bar means that the HRS was absent. The protein where each HRS maps is shown on the right. The HRS associations are based on their presence in the same HCV sample, deduced from the same or different amplicons; limitations for conclusions on HRS linkage in the same genome explained in the legend for Figure 2 apply also here.

A distribution of individual HRS among the twelve different DAA-based treatments undergone by the patients confirmed both the prevalence of HRS prior to the therapy and their maintenance (in most cases) following treatment, both regarding the percentage of patients with a given HRS, and the frequency of each HRS in the patient quasispecies (Figure 4). Therefore, HRSs at treatment failure are largely determined by their presence prior to DAA treatment.

**Figure 4.**
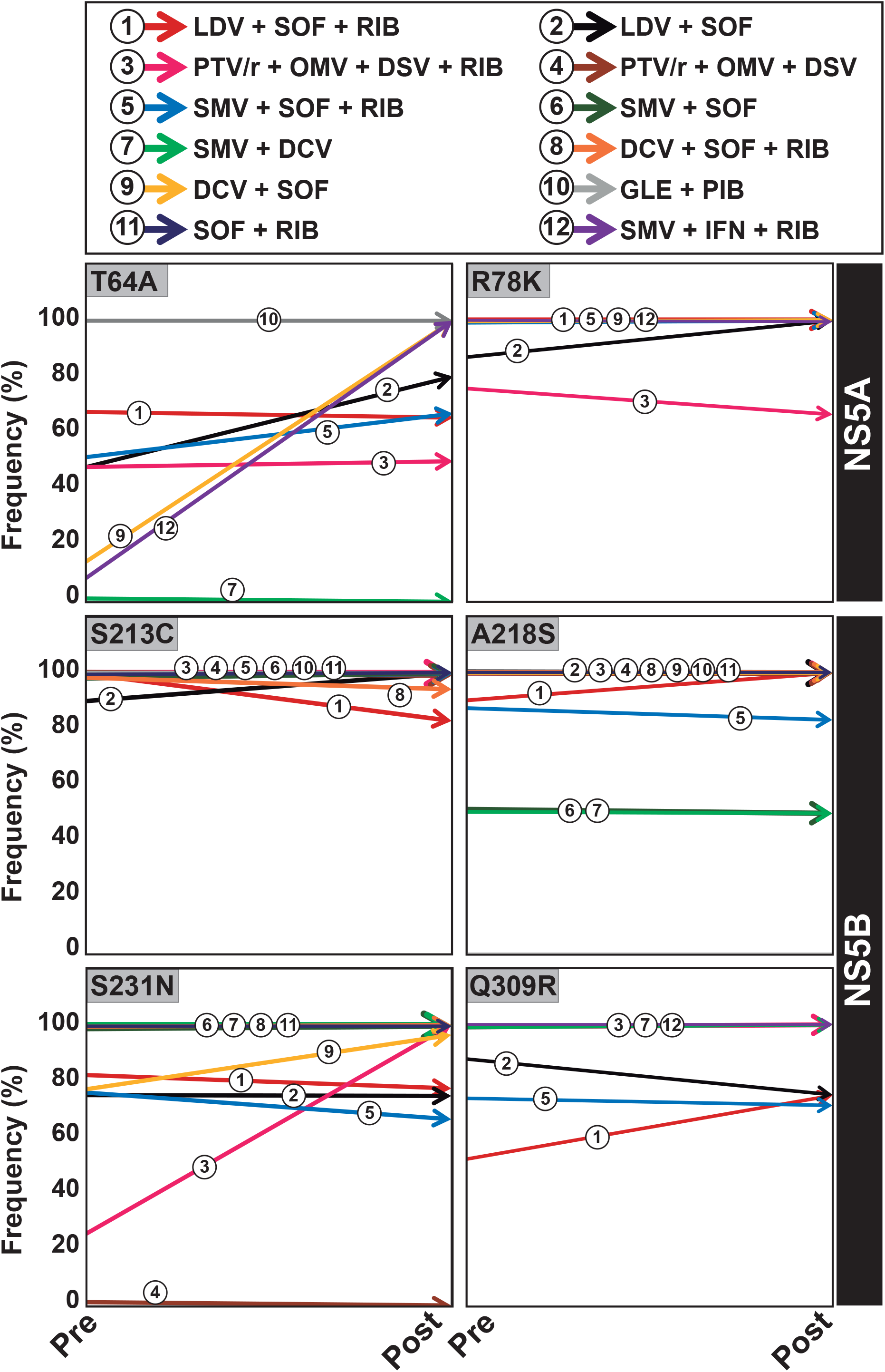
Comparison of individual HRSs in HCV in the basal (pre-treatment) and failure (post-treatment) samples according to the treatment. Frequency of each HRS in the viral quasispecies in the basal (pre-treatment) and failure (post-treatment) samples. For each HRS (given inside each panel) and treatment (top box), the frequency value for pre and post samples (arrows with color code in the top box) were calculated as an average of the values for the patients that underwent the indicated treatment. Drug abbreviations: LDV: ledipasvir; SOF: sofosbuvir; RIB: ribavirin; PTV/r: paritaprevir/ritonavir; OMV: ombitasvir; DSV: dasabuvir; SMV: simeprevir; DCV: daclatasvir; GLE: glecaprevir; PIB: pibrentasvir; IFN: interferon.

### Association of HRS with RAS

To evaluate the possible association between HRS and RAS in our cohort, we quantified the number of patients whose HCV contained both substitution categories in the virus sample obtained after treatment failure (Figure S2). In most cases there is a statistically significant association of HRSs and RASs in HCV from patients who failed therapy: 90% of patients were infected with HCV harboring at least one HRS combined with RAS whereas only 10% carried at least one HRS without RAS (p = 2.2×10^−16^; proportion test). Interestingly, combinations of the RAS L159F or C316N with the HRSs S213C or A218S are statistically significant as compared to combinations with other HRSs such as T64A, R78K, S231N and Q309R (Table S4). Also, combinations of the HRSs S213C+A218S and S213C+A218S+S231N were mainly associated with RAS L159F and C316N. Specifically, in all patients whose HCV contains RAS C316N, HRS A218S was also found.

### Tolerance of the substitution repertoire at treatment failure

To investigate possible differences between the amino acids that conform the HRS criterion, those involved in RAS, and other substitutions found at lower frequency in the same cohort following treatment failure, the acceptability of each substitution was quantified according to the PAM250 matrix (Figure S3). This value provides three levels of substitution acceptability based on amino acid structural resemblance and genetic inter-convertibility (PAM250<0, lower acceptability than expected, meaning a rare replacement; PAM250=0, acceptability as expected; PAM250>0 acceptability higher than expected) [34]. More than fifty percent (59±4.7%) of the total number of amino acid substitutions (HRS, RAS, and others) found at treatment failure display a PAM250 value higher than 0, indicating a high average acceptability of amino acid substitutions (Figure S3A). The most salient difference is that 19.3% of all substitutions belong to PAM250<0 while for RAS the proportion increases to 38.5% for the NS5B region (Figure S3A and S3B). In contrast, all HRS correspond to PAM250 ≥ 0 (Figure S3C).

To evaluate if a PAM250 acceptance category was associated with the percentage of patients harboring an amino acid substitution in each category, the frequency of patients in which any amino acid substitution occurred was divided in six categories: >20%, 15%-19.9%, 10%-14.9%, 5%-9.9%, 1%-4.9%, and 0.5%-0.9%. The amino acid substitutions in NS3, NS5A and NS5B that belong to a PAM250 category were distributed among the patient frequency groups (Figure S3D). The less accepted substitutions (those with PAM250<0) were mainly found in the low frequency groups of patients, with a difference that was statistically significant relative to the higher patient frequency groups (Table S5). These associations are expected from the fact that replacements that are not well tolerated tend to inflict a larger fitness cost upon the virus, thus attaining lower frequency among patients than those with high acceptance. Of note, the HRSs characterized in our study belong to well tolerated class of amino acid substitutions.

### Substitution tolerance and its relationship with residue conservation in data banks

We have recently described that amino acid conservation in the Los Alamos HCV databank (LANL) (https://hcv.lanl.gov/content/sequence/HCV/ToolsOutline.html) did not match low amino acid substitution frequency in HCV mutant spectra quantified both in cell culture and in HCV-infected patients [35]. In view of this unexpected result, it was interesting to determine the relationship between PAM250 values and the conservation range of amino acid positions according to the LANL alignment. The number of substitutions belonging to a defined PAM250 category and the total number of substitutions (comprising all PAM250 categories) was plotted as a function the degree of conservation of the amino acid at each position, calculated relative to the most abundant residue in the corresponding position of the LANL alignment (Figure 5 A-D). In all cases, the distribution followed a pattern which is very similar to that previously described for several HCV quasispecies at the nucleotide and amino acid level [35]. Only minor differences were noted in the distribution calculated for different PAM250 categories, and they did not reach statistical significance (p=0.7865, chi-square test with Monte Carlo correction).

**Figure 5.**
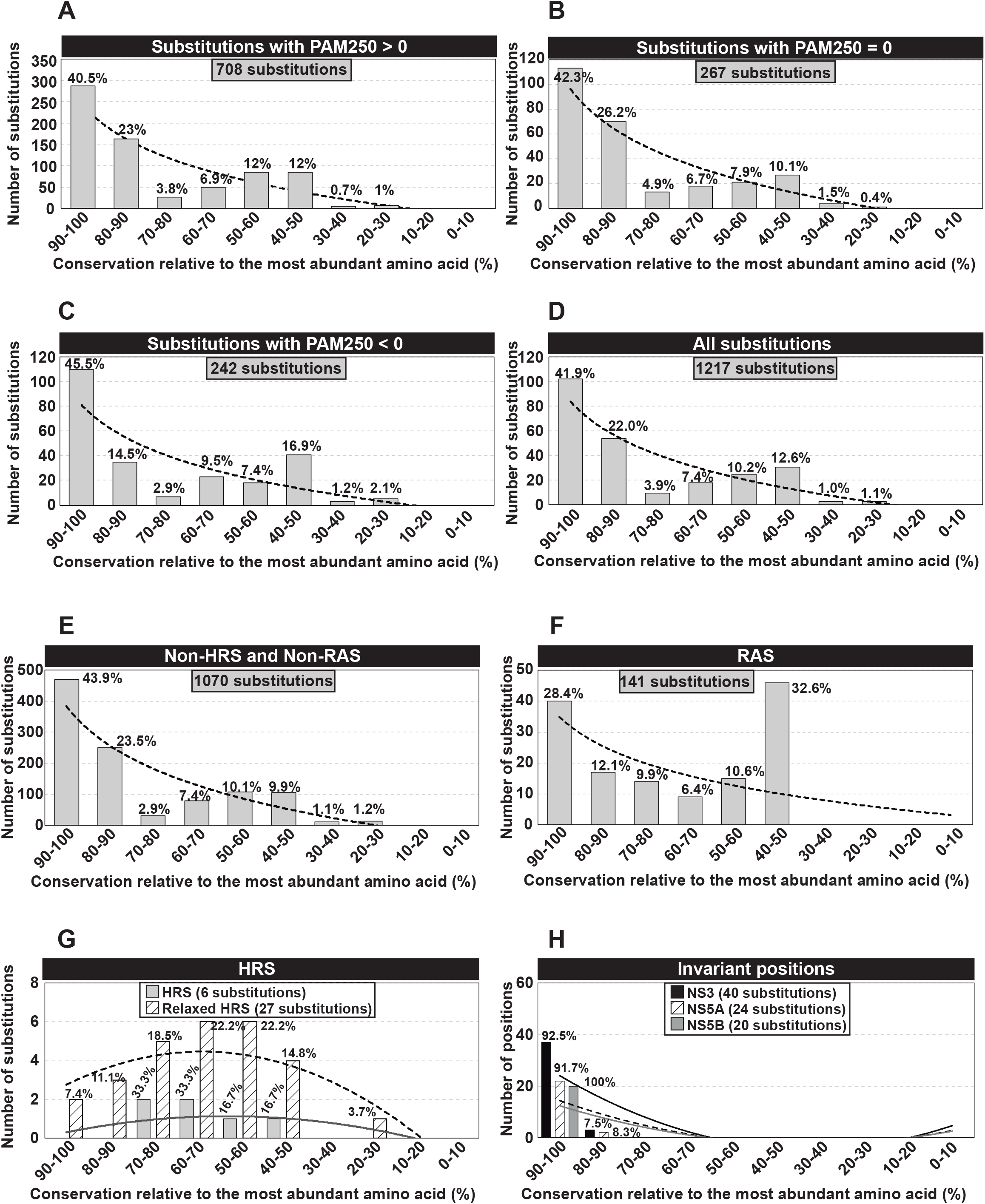
Number of amino acid substitutions in HCV from patients who failed therapy, distributed among conservation groups according to the LANL alignment. **(A)** Number of substitutions with PAM250>0 distributed among conservation groups, calculated relative to the most abundant amino acid at the corresponding position in the LANL alignment. Conservation groups are indicated in abscissa, and the number of substitutions in each group is given in ordinate (grey bars). The percentage of amino acid substitutions belonging to each category is indicated above each bar. The discontinuous line corresponds to function y = −112.8ln(x) + 241.16 (R^2^ = 0.8045). **(B)** Same as A but with amino acid substitutions with PAM250=0. The discontinuous line corresponds to function y = −46.22ln(x) + 96.506 (R^2^ = 0.8452). **(C)** Same as A but with substitutions with PAM250<0. The discontinuous line corresponds to function y = −37.85ln(x) + 81.366 (R^2^ = 0.6868). **(D)** Same as A but with the total number of amino acid substitutions. The discontinuous line corresponds to function y = −196.8ln(x) + 419.03 (R^2^ = 0.8035). **(E)** Number of amino acid substitutions, excluding HRSs and RASs, distributed among conservation groups, calculated relative to the most abundant amino acid at the corresponding position in the LANL alignment. The discontinuous line corresponds to function y = −182.9ln(x) + 383.26 (R^2^ = 0.8096). **(F)** Same as E but with RAS. The discontinuous line corresponds to function y = −13.81ln(x) + 34.955 (R^2^ = 0.366). **(G)** Same as E but with HRSs (present in more than 20% of patients), and relaxed HRSs (present in more than 10% of patients). The grey line corresponds to function y = −0.0606x^2^ + 0.5697x – 0.2 (R^2^ = 0.4242) for HRSs, and the discontinuous line corresponds to function y = −0.1742x^2^ + 1.4379x + 1.5 (R^2^ = 0.6459) for relaxed HRSs. **(H)** Same as E but with invariant positions in NS3, NS5A and NS5B. The black line corresponds to function y = 0.8636x^2^ – 11.645x + 34.8 (R^2^ = 0.6351) for NS3; the discontinuous line corresponds to the function y = 0.5152x^2^ – 6.9515x + 20.8 (R^2^ = 0.642) for NS5A; and the grey line corresponds to the function y = 0.4545x^2^ – 6.0909x + 18 (R^2^ = 0.5758) for NS5B. The complete list, location, statistical acceptability, and frequency among patients of all amino acid substitutions is given in Table S1.

The distribution pattern was also very similar when both HRS and RAS were excluded (Figure 5E). In contrast, the distribution of the 141 RAS substitutions differed in that 32.6% of the substitutions fell into the 40%-50% conservation category (p = 2.77 × 10^−14^; proportion test) (Figure 5F). The HRSs were spread among intermediate conservation categories (Figure 5G). A similar distribution was found when the cut-off level of percentage of patients with non-RAS substitutions was relaxed to 10% (Figure 5G). The previously described distribution of quasispecies mutations and amino acid substitutions among the LANL conservation groups denotes a higher tolerability of mutations at the quasispecies level than reflected in the consensus sequences that compose the LANL databank [35]. This implies that the HRSs described in our study are not constrained by a limitation of acceptability or by belonging to the most conserved amino acids according to the LANL.

Forty, twenty-four, and twenty positions in NS3-, NS5A- and NS5B-coding regions remained invariant, and were spread among the most conserved categories (80%-100%) (Figure 5H).

### Localization of HRS in the NS5A and NS5B structure

The three-dimensional structures of NS5A and NS5B proteins of HCV genotype 1b were used to localize the RAS and HRS positions [Protein Data Bank (http://www.wwpdb.org/), accession numbers 1ZH1and 5TRH for NS5A and NS5B, respectively] (Figure 6). In NS5A, the HRS positions 64 and 78 are located at the protein surface with side chains totally exposed to the solvent. Substitutions T64A and R78K can be easily accommodated without causing major distortions in structure (Figure 6A). Substitutions in position 93 represent one of the major antiviral resistance changes in HCV genotypes 1a and 1b [36]. As shown in Figure 6A, the NS5A position 93 can accommodate both Tyr and His side chains, maintaining similar neighboring interactions.

**Figure 6.**
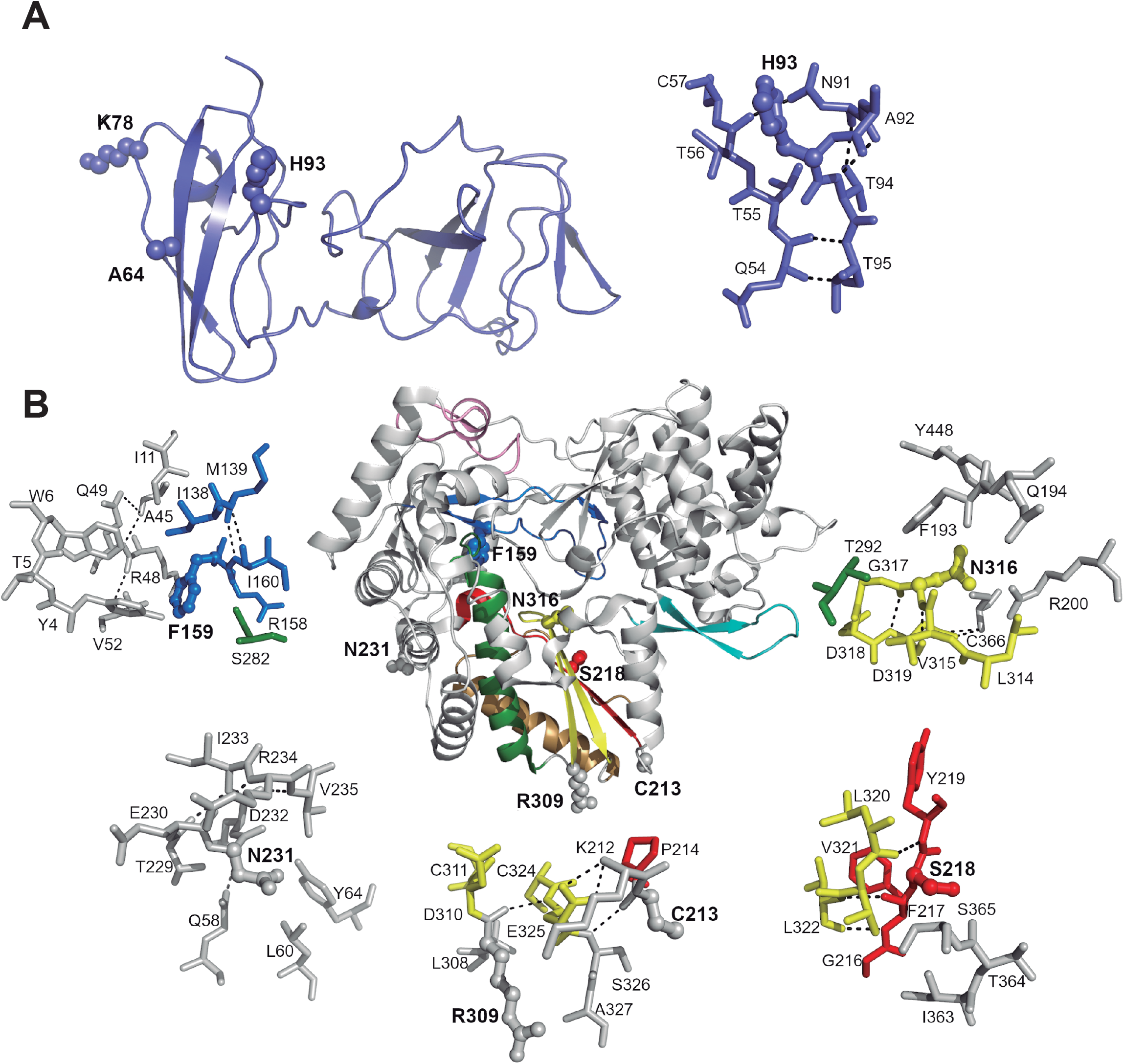
Positioning of amino acid substitutions, in NS5A and NS5B proteins, associated with treatment failure in HCV infection. **(A)** Cartoon representation of the NS5A protein structure (light blue) from HCV genotype 1b (PDB id 1ZH1), where the HRS T64A, R78K and RAS Y93H substitutions have been depicted as spheres (left panel). The right panel shows a close up view of the H93 substitution with surrounding residues in a 5 Å radius. **(B)** Cartoon representation of the HCV NS5B (RdRP) (genotype 1b; PDB id 5TRH) with the conserved structural motifs highlighted in different colors (A, red; B, green; C, yellow; D, sand; E, cyan; F, blue; G, pink). The side chains of residues with RAS (L159F, C316N) and HRS (S213C, A218S, S231N, Q309R) substitutions have been shown as spheres and labelled (central panel). Close up views of the substituted amino acids (ball and sticks and bold labels) with neighboring residues, in a 5 Å radius. The RAS substitution F159 is located within motif F of NS5B in close contact with a cluster of hydrophobic and aromatic residues (upper left panel). The RAS substitution N316 (top right panel) is in palm motif C, in a position previous to the catalytic site (G317D318D319). In contrast, HRS substitutions are located in highly exposed regions: in the fingers domain, N231(bottom left panel), at the base of the palm, C213 and R309 (bottom center panel), and in the β-strand that conforms the palm motif A, S218 (bottom right panel).

Similar to that observed in the NS5A structure, the amino acid positions 213, 231 and 309 in the NS5B polymerase, including most of HRS positions, are also solvent exposed at the protein surface, and replacements S213C, S231N and Q309R can be easily accommodated in the structure without distortions. Residues 213 and 309 are located at the base of the palm subdomain and amino acid 231 is in an exposed position in the polymerase fingers (Figure 6B). The calculated distances between these amino acid positions and the active site are 26.5 Å, 22 Å and 17.2 Å, respectively, for amino acids C213, R309 and N231. Considering that these three residues are far from the active site, it is not expected that such substitutions could affect the activity of the polymerase.

The HRS A218S is located in the palm subdomain, within the β-strand that conforms motif A (Figure 6B), and exposed on the surface of the NTP tunnel. It is also in close proximity to the catalytic Asp residues (at 6.5 Å distance of D220 in motif A and at 11 Å of D318 of motif C). S218 is also close to the RAS mutation N316 (11.4 Å). However, neither the A218S nor C316N substitutions appear to distort the structure of polymerase catalytic site. The potential impact of the C316N mutation on efficacy of the antiviral drug sofosbuvir in patients infected with HCV genotype1b has been previously evaluated. The authors suggested that the bulkier N316 side chain would partially block the access of the nucleotide analog to the polymerase active site by inducing steric hindrance with the additional 2’Me and 2’F groups of sofosbuvir compared to natural nucleotides [37].

Finally, RAS L159F is located in the fingers motif F, forming part of the template channel (Figure 6B). However, its side chain is not oriented towards the channel but packed towards the polymerase interior, participating in hydrophobic cluster. The replacement of L159 by the bulkier F side chain would add new interactions to the cluster, though causing only minimal distortions (Figure 6B).

### A comparison of HRS frequency between a cohort of HCV-patients with treatment failure and a cohort with sustained viral response

To investigate if HRSs might have predictive value regarding DAA-based treatment outcome, we compared the presence of each HRS in the basal samples of our cohort [9] whose outcome was treatment failure, and in basal samples of another cohort [38] whose outcome was sustained virological response (SVR). The results (Figure 7) indicate statistically significant differences in the percentage of patients who carried some (but not all) HRSs (Table S6). The most striking difference was the absence of T64A in HCV genotype G1a in those patients who achieved SVR. The observed differences open the possibility that some HRSs may assist in predicting treatment response, a suggestion that must await identification of additional HRSs in other cohorts.

**Figure 7.**
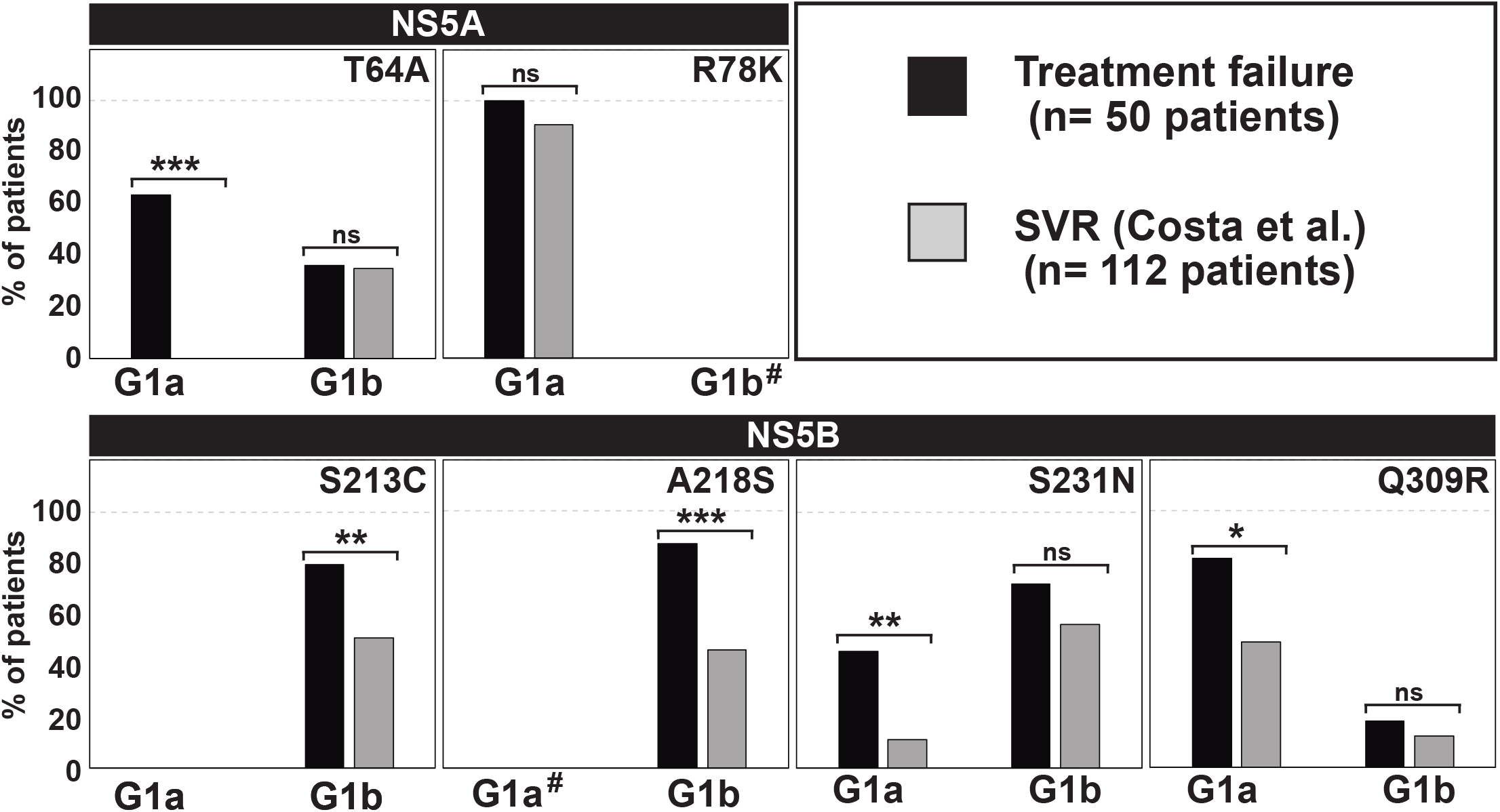
A comparison of HRS occurrence in basal samples of our cohort and in a cohort of patients who achieved sustained virological response. The presence of an HRS (indicated in each panel) was examined in the 50 basal (prior to treatment) samples of our patient cohort (8) whose outcome was treatment failure (black bars), and 112 basal samples from another patient cohort (34) whose outcome was sustained virological response (SVR) (grey bars). The HCV genotype is given in abscissa, and the percentage of patients carrying a HRS is written in ordinate. The statistical significance of the difference in the frequency of HRS between the two cohorts is indicated (ns=not significant; * p<0.05; ** p<0.01; *** p<0.001; proportion test). #means that the HRS R78K in NS5A and A218S in NS5B cannot be considered because the reference amino acid for G1b is a K and for G1a an S, respectively.

## Discussion

HCV replicating in infected patients displays the complex quasispecies dynamics expected from low-fidelity replication in any environment, independently of external perturbations [39,40]. New mutations arise and vary in frequency as a function of time, due to selective forces and random sampling events. Only a subset of all mutations become dominant in response to antiviral agents. Such a subset conforms the list of RAS that, in the case of HCV, is periodically updated with the aims of interpreting antiviral intervention failures and to aid in treatment planning [6]. However, and important, not all amino acid substitutions that vary in frequency during intra-host HCV evolution, and that increase their frequency in the quasispecies, need be the result of direct selection by antiviral agents. Several reports have described amino acid substitutions in cohorts of treated and untreated HCV-infected patients that are not *bona fide* RAS, and whose contribution to treatment failures is uncertain [37,41–44].

We have examined this open question with a large cohort of 220 HCV-infected patients that failed DAA-based therapies, since 25 patients did not exhibit any known RAS after treatment failure [9]. Using UDS information of the 220 HCV-infected patients, we have characterized a new class of amino acid substitutions that we have termed HRS. We examined the proportion of patients in whom they are present, their frequency within the mutant spectrum of the quasispecies, and their statistical acceptability based on parameters employed in studies of protein evolution. An HRS differs from a RAS in the following features: (i) they are not listed as RAS in current RAS catalogues [6,31,32]; (ii) they can be found in basal samples and remain dominant in patients undergoing different DAA treatments; (iii) they correspond to well accepted substitutions according to the PAM250 replacement matrix, and (iv) they belong to intermediate conservation categories according to the conservation range of amino acid positions in the LANL alignment.

Some of the HRSs identified in our cohort are dominant for a specific HCV subtype but not for others (i.e. in position 78 in NS5A, the wt (reference) amino acid is a R for G1a but K for G1b; in position 218 in NS5B, the wt (reference) amino acid is S for G1a but A for G1b) [also found in [37,38]]. Additionally, while HRS combinations in basal samples amounted to 62.5% of patients infected with HCV G1a, they represented only 12.5 % for infections with G1b. These differences argue in favor of an influence of the HCV genetic background in HRS occurrence and prominence. HRS acceptability is also consistent with the limited perturbations predicted to inflict on the proteins harboring them, according to modeling of the effect of the relevant substitutions on the three-dimensional structure of the proteins. The fact that 34.5% of patients carrying one or more HRSs were neither treated nor failed to drugs whose target are the proteins where the HRSs were located, reinforces the lack of association between HRS presence and a specific treatment.

Some of the HRS that we have characterized, have been also reported in other cohorts, with no evidence of them being RAS. Uchida *et al.* identified A218S+C316N in the G1b viral population that became dominant upon failure to LDV+SOF, as well as in SOF-naïve patients [37]. We have also identified A218S in patients never subjected to SOF treatment, despite evidence that this substitution may jeopardize the access of SOF-triphosphate to the catalytic site of NS5B [37]. Bellocchi *et al.* (2019) identified K78R and T64A in NS5A and C213S, S231N, N231S and A218S in HCV G1b-infected patients naïve to DAAs [43]. T64S in NS5A was listed as a secondary substitution accompanying P58S and Y93H in HCV G1b-infected patients who failed DCV therapy, with a resistance level of EC_50_ <1 nmol/l [44].

The observations regarding Q309R are worth commenting. While this substitution has been previously associated with ribavirin (RIB) resistance [45,46], a direct specific involvement in RIB resistance is not obvious. For HCV G1b-infected patients, Kim *et al.* found Q309R at high frequency in the quasispecies of treatment-naïve patients [42], and Jiang et al. described it at lower frequency in G1b-infected patients, prior to treatment [47]. In our cohort, Q309R was present in the basal samples of several patients. Although we cannot exclude that some patients had undergone a prior pegIFN-α+RIB treatment, or that they were infected with virus from patients that had undergone RIB-containing therapies, the frequency of genomes with Q309R remained high during DAA therapy, independently of their including RIB.

Discrepancies between clinical observations and results of the effect of mutations using replicon systems [37], added to the multiple genetic backgrounds in which a specific amino acid substitution (alone or in combination) should be tested *in vitro*, renders very difficult a definitive assignment of a substitution to the HRS category and its total exclusion from any RAS activity.

The possible origin of HRS as a result of some selective constraint that acted during the prior evolutionary history of the virus cannot be excluded, but identification of the potential selective agent is challenging. For example, none of the HRS we have characterized maps within conserved T cell epitopes predicted in NS5A and NS5B by bioinformatic procedures [48], suggesting that the origin of HRS is unrelated to escape from cellular immune responses. Probably, we have identified only a minimum subset of the constraints to which viruses are subjected in their natural environments. One possibility is that HRS may be prompted by their favoring viral fitness irrespectively of being or not together with a RAS in the same genome. Fitness has multiple survival-enhancing effects on evolving viral populations [49]. A possible survival value of HRS should be more noticeable in the presence of those RAS that inflict a fitness cost. Moreover, using HCV infection of human hepatoma cells in culture, we have documented that fitness *per se* (in absence of any RAS, ascertained by different procedures) is a determinant of HCV multidrug resistance, including IFN-α, DAAs, cyclosporine A (that targets a cellular protein), and the mutagenic analogues favipiravir and RIB [27–29]. In view of these results, it is tempting to consider that the HRS class of substitutions may play a fitness-enhancing role. They would be the counterpart in infected patients of the multiple mutations scattered along the HCV genome that have been associated with fitness increase of HCV in cell culture [50].

Regarding the diagnostic relevance of HRS, if our findings are confirmed with additional patient cohorts, a baseline identification of HRS may provide information to be added to other predictors of treatment outcome, be them RAS presence, or quasispecies complexity [51]. As an example, Mawatari et al. observed that in a cohort of G1b-infected patients, when A218S and C316N were absent, SVR was achieved in all cases [41]. Sequencing of basal samples is a recommendation for treatment planning. Therefore, the information on HRS that will be gathered during sequencing should be relevant not only to help predicting treatment outcomes, but also to further understand HCV population dynamics which appears much more complex than thought prior to introduction of deep sequencing.

## Materials and Methods

### Experimental data on HCV-infected patients

DNA amplifications from viral RNA was performed using subtype-specific oligonucleotides previously described. Amplification and PCR-mediated recombination errors, and reproducibility the same or different sequencing platforms were controlled experimentally and bioinformatically [33,52]. Deep sequencing procedures, patient clinical data and HCV sequences from infected patients have been previously described [9]. The cut-off frequency of amino acid substitution detection was 1% [33]. Similar procedures were used for the amplification and sequencing of pre-treatment (basal) and post-treatment samples.

### Sequences from the Los Alamos data base

The sequences were retrieved from LANL following previously described procedures [33]. Inclusion criteria were that the sequences had been confirmed, with no evidence of their being recombinants, and that they corresponded to full-length (or near-full length) genomes (without large insertions or deletions). Their HCV genotype / subtype distribution is: 553 sequences of genotype G1a; 427 of G1b; 3 of G1c; 33 of G2a; 81 of G2b; 7 of G2c; 5 of G2j; 3 of G2k, although no distinction between HCV subtypes was made for the calculation of the conservation range of individual residues. Alignments were performed using the program BioEdit version 7.0.9.0.

### Statistics

The statistical significance of differences in the distribution of variable sites among conservation groups was calculated with the Pearson’s chi-square test using software R version 3.6.2, with Monte Carlo correction (based on 2000 replicates). Sample sizes are given for each comparison. The proportion test using software R version 3.6.2 was used for multiple determinations.

### Sequence accession numbers and data availability

The reference accession numbers of sequences retrieved from LANL used to determine conservation groups are given in [35]. Accession numbers for HCV samples included in the patient cohort are SAMN08741670 to SAMN08741673 [33]. Amino acid replacements in HCV from infected patients have been compiled in Table S1.

## Acknowledgments

The work at CBMSO was supported by grants SAF2014-52400-R from Ministerio de Economía y Competitividad (MINECO), SAF2017-87846-R, BFU2017-91384-EXP from Ministerio de Ciencia, Innovación y Universidades (MICIU), PI18/00210 from Instituto de Salud Carlos III, S2013/ABI-2906, (PLATESA from Comunidad de Madrid/FEDER) and S2018/BAA-4370 (PLATESA2 from Comunidad de Madrid/FEDER). C.P. is supported by the Miguel Servet program of the Instituto de Salud Carlos III (CP14/00121 and CPII19/00001) cofinanced by the European Regional Development Fund (ERDF). CIBERehd (Centro de Investigación en Red de Enfermedades Hepáticas y Digestivas) is funded by Instituto de Salud Carlos III. Institutional grants from the Fundación Ramón Areces and Banco Santander to the CBMSO are also acknowledged. The team at CBMSO belongs to the Global Virus Network (GVN). The work in Barcelona was supported by Instituto de Salud Carlos III, cofinanced by the European Regional Development Fund (ERDF) grant number PI19/00301 and by the Centro para el Desarrollo Tecnológico Industrial (CDTI) from the MICIU, grant number IDI-20151125. Work at CAB was supported by MINECO grant BIO2016-79618R and PID2019-104903RB-I00 (funded by EU under the FEDER program) and by the Spanish State research agency (AEI) through Project number MDM-2017-0737 (Unidad de Excelencia “María de Maeztu”-Centro de Astrobiología (CSIC-INTA). Work at IBMB was supported by MICIN grant BIO2017-83906-P (funded by EU under the FEDER program). C. G.-C. is supported by predoctoral contract PRE2018-083422 from MICIU. B. M.-G. is supported by predoctoral contract PFIS FI19/00119 from Instituto de Salud Carlos III (Ministerio de Sanidad y Consumo) cofinanced by Fondo Social Europeo (FSE).

