## Supplemental Material for "Amino acid substitutions associated with treatment failure of hepatitis C virus infection"

**Table S1. Repertoire of amino acid substitutions in NS3-, NS5A- and NS5B-coding regions identified in HCV infected patients[a].**

| | Region | Substitution[b] | PAM250[c] | Number of patients | % of patients |
|---|---|---|---|---|---|
| > 20% of patients | NS5A | T64A | 1 | 54 | 24.5 |
| | | R78K | 3 | 45 | 20.5 |
| | | Y93H* | 0 | 74 | 33.6 |
| | NS5B | L159F* | 2 | 46 | 20.9 |
| | | S213C | 0 | 72 | 32.7 |
| | | A218S | 1 | 70 | 31.8 |
| | | S231N | 1 | 57 | 25.9 |
| | | Q309R | 1 | 46 | 20.9 |
| | | C316N* | -4 | 54 | 24.5 |
| 15-19.9% of patients | NS3 | V48I | 4 | 33 | 15.0 |
| | | Y56F[+] | 7 | 40 | 18.2 |
| | | A150V | 0 | 40 | 18.2 |
| | | V170I[+] | 4 | 37 | 16.8 |
| | NS5A | L37F | 2 | 43 | 19.5 |
| | | Q123R | 1 | 35 | 15.9 |
| | | T135A | 1 | 40 | 18.2 |
| | NS5B | K124E | 0 | 38 | 17.3 |
| | | P189S | 1 | 35 | 15.9 |
| 10-14.9% of patients | NS3 | T46S | 1 | 27 | 12.3 |
| | | I71V | 4 | 22 | 10.0 |
| | | T72I | 0 | 24 | 10.9 |
| | | S147L | -3 | 28 | 12.7 |
| | | D168V* | -2 | 27 | 12.3 |
| | | N174S | 1 | 31 | 14.1 |
| | NS5A | L31M* | 4 | 26 | 11.8 |
| | | R44K | 3 | 29 | 13.2 |
| | | R48K | 3 | 24 | 10.9 |
| | | Q54H | 3 | 28 | 12.7 |
| | | K78R | 3 | 31 | 14.1 |
| | | H85Y | 0 | 22 | 10.0 |
| | | V138L | 2 | 26 | 11.8 |
| | NS5B | V150A | 0 | 23 | 10.5 |
| | | R254K | 3 | 27 | 12.3 |
| | | S300T | 1 | 26 | 11.8 |
| 5-9.9% of patients | NS3 | V33I | 4 | 12 | 5.5 |
| | | A40T | 1 | 13 | 5.9 |
| | | P67S | 1 | 18 | 8.2 |
| | | A67V | 0 | 16 | 7.3 |
| | | Q86P | 0 | 12 | 5.5 |
| | | P89S | 1 | 14 | 6.4 |
| | | S93P | 1 | 19 | 8.6 |
| | | T98A | 1 | 21 | 9.5 |
| | | S102P | 1 | 12 | 5.5 |
| | | S125G | 1 | 16 | 7.3 |
| | | V132I[+] | 4 | 19 | 8.6 |
| | | S147M | -2 | 13 | 5.9 |
| | | A166T | 1 | 11 | 5.0 |
| | NS5A | R30Q* | 1 | 13 | 5.9 |

| | | | | | |
|---|---|---|---|---|---|
| | | Q30R* | 1 | 19 | 8.6 |
| | | L31I* | 2 | 11 | 5.0 |
| | | F37L | 2 | 18 | 8.2 |
| | | K44R | 3 | 14 | 6.4 |
| | | V46A | 0 | 14 | 6.4 |
| | | R48Q | 1 | 13 | 5.9 |
| | | T56I | 0 | 16 | 7.3 |
| | | T58P* | 0 | 21 | 9.5 |
| | | T64S | 1 | 14 | 6.4 |
| | | A75V | 0 | 12 | 5.5 |
| | | T79A | 1 | 13 | 5.9 |
| | | S85N | 1 | 11 | 5.0 |
| | | S103P | 1 | 20 | 9.1 |
| | | S107T | 1 | 11 | 5.0 |
| | | V121I | 4 | 11 | 5.0 |
| | | I130V | 4 | 11 | 5.0 |
| | | V138I | 4 | 14 | 6.4 |
| | | A147P | 1 | 16 | 7.3 |
| | NS5B | K124Q | 1 | 12 | 5.5 |
| | | E124- | - | 16 | 7.3 |
| | | S130N | 1 | 18 | 8.2 |
| | | E131D | 3 | 15 | 6.8 |
| | | V147I | 4 | 13 | 5.9 |
| | | T181N | 0 | 20 | 9.1 |
| | | L184V | 2 | 17 | 7.7 |
| | | G198A | 1 | 20 | 9.1 |
| | | A207T | 1 | 20 | 9.1 |
| | | K212R | 3 | 11 | 5.0 |
| | | R250K | 3 | 11 | 5.0 |
| | | I262V | 4 | 15 | 6.8 |
| | | R270K | 3 | 16 | 7.3 |
| | | N273S | 1 | 12 | 5.5 |
| | | S300A | 1 | 12 | 5.5 |
| | | K304R | 3 | 16 | 7.3 |
| **1-4.9% of patients** | NS3 | V35I | 4 | 3 | 1.4 |
| | | A39T | 1 | 7 | 3.2 |
| | | S42T | 1 | 5 | 2.3 |
| | | F43L* | 2 | 5 | 2.3 |
| | | I48V | 4 | 8 | 3.6 |
| | | V48L | 2 | 3 | 1.4 |
| | | V51A | 0 | 8 | 3.6 |
| | | V55A* | 0 | 3 | 1.4 |
| | | Y56H* | 0 | 9 | 4.1 |
| | | S61A | 1 | 4 | 1.8 |
| | | S61T | 1 | 6 | 2.7 |
| | | R62K | 3 | 3 | 1.4 |
| | | T63A | 1 | 4 | 1.8 |
| | | T63S | 1 | 3 | 1.4 |
| | | I64L | 2 | 3 | 1.4 |
| | | G66S | 1 | 3 | 1.4 |
| | | K68N | 1 | 5 | 2.3 |
| | | H69R | 2 | 4 | 1.8 |
| | | T72V | 0 | 3 | 1.4 |

| | | | | | |
|---|---|---|---|---|---|
| | | V78I | 4 | 3 | 1.4 |
| | | Q80R* | 1 | 9 | 4.1 |
| | | Q80K* | 1 | 10 | 4.5 |
| | | Q80L* | -2 | 4 | 1.8 |
| | | D81G | 1 | 3 | 1.4 |
| | | V83I | 4 | 5 | 2.3 |
| | | P88L | -3 | 3 | 1.4 |
| | | Q89P | 0 | 7 | 3.2 |
| | | A91T | 1 | 3 | 1.4 |
| | | A91S | 1 | 3 | 1.4 |
| | | A91V | 0 | 3 | 1.4 |
| | | K92R | 3 | 3 | 1.4 |
| | | L94M | 4 | 4 | 1.8 |
| | | T95A | 1 | 4 | 1.8 |
| | | T95P | 0 | 6 | 2.7 |
| | | P96S | 1 | 4 | 1.8 |
| | | T98S | 1 | 5 | 2.3 |
| | | T98P | 0 | 9 | 4.1 |
| | | S101P | 1 | 9 | 4.1 |
| | | S101R | 0 | 4 | 1.8 |
| | | A102V | 0 | 7 | 3.2 |
| | | S102A | 1 | 3 | 1.4 |
| | | D103A | 0 | 5 | 2.3 |
| | | D103G | 1 | 3 | 1.4 |
| | | V107I* | 4 | 6 | 2.7 |
| | | R109G | -3 | 6 | 2.7 |
| | | H110P | 0 | 4 | 1.8 |
| | | D110E | 3 | 6 | 2.7 |
| | | D112A | 0 | 4 | 1.8 |
| | | V113I | 4 | 4 | 1.8 |
| | | I114V | 4 | 6 | 2.7 |
| | | I114L | 2 | 9 | 4.1 |
| | | R117H | 2 | 7 | 3.2 |
| | | S122T* | 1 | 10 | 4.5 |
| | | S122N* | 1 | 5 | 2.3 |
| | | S122G* | 1 | 5 | 2.3 |
| | | T122A$^{+}$ | 1 | 3 | 1.4 |
| | | A133S | 1 | 8 | 3.6 |
| | | S134T | 1 | 3 | 1.4 |
| | | P146S | 1 | 3 | 1.4 |
| | | S147A | 1 | 4 | 1.8 |
| | | A151T | 1 | 3 | 1.4 |
| | | V151A | 0 | 3 | 1.4 |
| | | I153V | 4 | 10 | 4.5 |
| | | R155K* | 3 | 8 | 3.6 |
| | | R155Q* | 1 | 6 | 2.7 |
| | | K165E | 0 | 3 | 1.4 |
| | | K165R | 3 | 3 | 1.4 |
| | | A166S | 1 | 5 | 2.3 |
| | | D168E* | 3 | 10 | 4.5 |
| | | D168A* | 0 | 6 | 2.7 |
| | | I170V* | 4 | 6 | 2.7 |
| | | N174G | 0 | 8 | 3.6 |

| | | | | | |
|---|---|---|---|---|---|
| | | T174A | 1 | 4 | 1.8 |
| | | T174I | 0 | 3 | 1.4 |
| | | S174A | 1 | 3 | 1.4 |
| | | S174F | -3 | 3 | 1.4 |
| | | M175L* | 4 | 5 | 2.3 |
| | | S176N | 1 | 8 | 3.6 |
| | | T177A | 1 | 4 | 1.8 |
| | | M179V | 2 | 3 | 1.4 |
| | | M179T | -1 | 3 | 1.4 |
| | | M179I | 2 | 5 | 2.3 |
| | | A179V | 0 | 7 | 3.2 |
| | | A179T | 1 | 6 | 2.7 |
| | NS5A | Q24R+ | 1 | 3 | 1.4 |
| | | A25T | 1 | 3 | 1.4 |
| | | S25A | 1 | 5 | 2.3 |
| | | S25T | 1 | 10 | 4.5 |
| | | L28S* | -3 | 3 | 1.4 |
| | | L28M* | 4 | 6 | 2.7 |
| | | L28V* | 2 | 7 | 3.2 |
| | | M28V* | 2 | 6 | 2.7 |
| | | M28T* | -1 | 3 | 1.4 |
| | | A30K* | -1 | 6 | 2.7 |
| | | A30S* | 1 | 4 | 1.8 |
| | | A30V+ | 0 | 3 | 1.4 |
| | | Q30E* | 2 | 3 | 1.4 |
| | | Q30K* | 1 | 4 | 1.8 |
| | | Q30H* | 3 | 3 | 1.4 |
| | | L31V* | 2 | 7 | 3.2 |
| | | M31I+ | 2 | 7 | 3.2 |
| | | M31V+ | 2 | 5 | 2.3 |
| | | I34V | 4 | 10 | 4.5 |
| | | V34I | 4 | 9 | 4.1 |
| | | V34L | 2 | 6 | 2.7 |
| | | F36L | 2 | 4 | 1.8 |
| | | I37V | 4 | 3 | 1.4 |
| | | L37I | 2 | 9 | 4.1 |
| | | R41K | 3 | 3 | 1.4 |
| | | V46I | 4 | 5 | 2.3 |
| | | I52V | 4 | 8 | 3.6 |
| | | M53T | -1 | 3 | 1.4 |
| | | H54Y | 0 | 3 | 1.4 |
| | | H54R | 2 | 3 | 1.4 |
| | | S54Y | -3 | 3 | 1.4 |
| | | Q54Y | -4 | 7 | 3.2 |
| | | T55A | 1 | 8 | 3.6 |
| | | T56A | 1 | 6 | 2.7 |
| | | P58S* | 1 | 7 | 3.2 |
| | | H58P+ | 0 | 3 | 1.4 |
| | | S62T+ | 1 | 9 | 4.1 |
| | | S62L+ | -3 | 4 | 1.8 |
| | | S62P+ | 1 | 3 | 1.4 |
| | | Q62E+ | 2 | 9 | 4.1 |
| | | I63V | 4 | 3 | 1.4 |

| | | | | | |
|---|---|---|---|---|---|
| | | K68R | 3 | 6 | 2.7 |
| | | N69T | 0 | 3 | 1.4 |
| | | S71P | 1 | 3 | 1.4 |
| | | R73K | 3 | 5 | 2.3 |
| | | I74V | 4 | 4 | 1.8 |
| | | V75I | 4 | 5 | 2.3 |
| | | V75A | 0 | 6 | 2.7 |
| | | T79K | 0 | 4 | 1.8 |
| | | M83T | -1 | 4 | 1.8 |
| | | M83V | 2 | 4 | 1.8 |
| | | T83M | -1 | 9 | 4.1 |
| | | S85G | 1 | 4 | 1.8 |
| | | T87A | 1 | 8 | 3.6 |
| | | I90V | 4 | 5 | 2.3 |
| | | A92T* | 1 | 4 | 1.8 |
| | | Y93S* | -3 | 3 | 1.4 |
| | | Y93C* | 0 | 4 | 1.8 |
| | | T95A | 1 | 3 | 1.4 |
| | | P97S | 1 | 4 | 1.8 |
| | | S98G | 1 | 10 | 4.5 |
| | | T99I | 0 | 3 | 1.4 |
| | | T99N | 0 | 3 | 1.4 |
| | | I101V | 4 | 6 | 2.7 |
| | | S101P | 1 | 7 | 3.2 |
| | | S103A | 1 | 4 | 1.8 |
| | | N105S | 1 | 4 | 1.8 |
| | | N105D | 2 | 4 | 1.8 |
| | | T107K | 0 | 9 | 4.1 |
| | | T107A | 1 | 4 | 1.8 |
| | | R108T | -1 | 3 | 1.4 |
| | | R108K | 3 | 9 | 4.1 |
| | | S114A | 1 | 3 | 1.4 |
| | | A114S | 1 | 4 | 1.8 |
| | | A114T | 1 | 3 | 1.4 |
| | | A115T | 1 | 4 | 1.8 |
| | | N116S | 1 | 8 | 3.6 |
| | | N116D | 2 | 7 | 3.2 |
| | | E117D | 3 | 3 | 1.4 |
| | | I121V | 4 | 8 | 3.6 |
| | | T122A | 1 | 3 | 1.4 |
| | | T122R | -1 | 4 | 1.8 |
| | | R123Q | 1 | 6 | 2.7 |
| | | D126E | 3 | 9 | 4.1 |
| | | E126D | 3 | 5 | 2.3 |
| | | F127L | 2 | 3 | 1.4 |
| | | F127S | -3 | 6 | 2.7 |
| | | M133T | -1 | 3 | 1.4 |
| | | M133V | 2 | 6 | 2.7 |
| | | M133I | 2 | 9 | 4.1 |
| | | T135V | 0 | 4 | 1.8 |
| | | E137G | 0 | 7 | 3.2 |
| | | N137D | 2 | 3 | 1.4 |
| | | I63V | 4 | 3 | 1.4 |

| | | | | | |
|---|---|---|---|---|---|
| | | K68R | 3 | 6 | 2.7 |
| | | A146T | 1 | 5 | 2.3 |
| | NS5B | K124R | 3 | 9 | 4.1 |
| | | K124D | 0 | 3 | 1.4 |
| | | K124G | -2 | 3 | 1.4 |
| | | K124M | 0 | 6 | 2.7 |
| | | E124K | 0 | 7 | 3.2 |
| | | E124G | 0 | 3 | 1.4 |
| | | E124Q | 2 | 4 | 1.8 |
| | | E128A | 0 | 3 | 1.4 |
| | | E128D | 3 | 4 | 1.8 |
| | | D129G | 1 | 3 | 1.4 |
| | | S130T | 1 | 3 | 1.4 |
| | | T130N | 0 | 8 | 3.6 |
| | | T130S | 1 | 7 | 3.2 |
| | | V131E | -2 | 4 | 1.8 |
| | | V131A | 0 | 7 | 3.2 |
| | | E131V | -2 | 10 | 4.5 |
| | | T131I | 0 | 8 | 3.6 |
| | | T132A | 1 | 3 | 1.4 |
| | | D135G | 1 | 3 | 1.4 |
| | | I138V | 4 | 5 | 2.3 |
| | | M139T | -1 | 3 | 1.4 |
| | | K141R | 3 | 3 | 1.4 |
| | | N142S | 1 | 10 | 4.5 |
| | | E143G | 0 | 4 | 1.8 |
| | | F145L | 2 | 5 | 2.3 |
| | | F145S | -3 | 5 | 2.3 |
| | | V147A | 0 | 5 | 2.3 |
| | | Q148E | 2 | 3 | 1.4 |
| | | Q148A | 0 | 6 | 2.7 |
| | | D148N | 2 | 3 | 1.4 |
| | | P149L | -3 | 4 | 1.8 |
| | | A150T | 1 | 3 | 1.4 |
| | | V150I | 4 | 3 | 1.4 |
| | | V150T | 0 | 9 | 4.1 |
| | | K151R | 3 | 6 | 2.7 |
| | | A156P | 1 | 3 | 1.4 |
| | | F162L | 2 | 5 | 2.3 |
| | | F162Y | 7 | 8 | 3.6 |
| | | V167A | 0 | 4 | 1.8 |
| | | V169I | 4 | 3 | 1.4 |
| | | C170R | -4 | 3 | 1.4 |
| | | K172E | 0 | 4 | 1.8 |
| | | D177G | 1 | 3 | 1.4 |
| | | D177N | 2 | 4 | 1.8 |
| | | S180K | 0 | 4 | 1.8 |
| | | K180R | 3 | 3 | 1.4 |
| | | T181K | 0 | 3 | 1.4 |
| | | T181S | 1 | 3 | 1.4 |
| | | L184P | -3 | 3 | 1.4 |
| | | L184Q | -2 | 10 | 4.5 |
| | | I184T | 0 | 5 | 2.3 |

| | | | | | |
|---|---|---|---|---|---|
| | | E185A | 0 | 5 | 2.3 |
| | | E185G | 0 | 3 | 1.4 |
| | | S189N | 1 | 3 | 1.4 |
| | | S189A | 1 | 3 | 1.4 |
| | | S189P | 1 | 6 | 2.7 |
| | | S190A | 1 | 4 | 1.8 |
| | | Y195H | 0 | 3 | 1.4 |
| | | S196P | 1 | 5 | 2.3 |
| | | S196T | 1 | 4 | 1.8 |
| | | G198S | 1 | 8 | 3.6 |
| | | Q199R | 1 | 5 | 2.3 |
| | | V201A | 0 | 3 | 1.4 |
| | | E202D | 3 | 8 | 3.6 |
| | | E202G | 0 | 4 | 1.8 |
| | | V205A | 0 | 4 | 1.8 |
| | | Q206R | 1 | 5 | 2.3 |
| | | K206Q | 1 | 7 | 3.2 |
| | | K206E | 0 | 9 | 4.1 |
| | | N206S | 1 | 6 | 2.7 |
| | | N206Q | 1 | 3 | 1.4 |
| | | N206D | 2 | 5 | 2.3 |
| | | N206K | 1 | 8 | 3.6 |
| | | K209R | 3 | 5 | 2.3 |
| | | K211E | 0 | 3 | 1.4 |
| | | K211R | 3 | 4 | 1.8 |
| | | K212E | 0 | 3 | 1.4 |
| | | S213T | 1 | 9 | 4.1 |
| | | S213N | 1 | 4 | 1.8 |
| | | M215V | 2 | 3 | 1.4 |
| | | F224L | 2 | 4 | 1.8 |
| | | F224S | -3 | 3 | 1.4 |
| | | T227A | 1 | 3 | 1.4 |
| | | V228A | 0 | 7 | 3.2 |
| | | V228I | 4 | 4 | 1.8 |
| | | E230G | 0 | 3 | 1.4 |
| | | S231G | 1 | 5 | 2.3 |
| | | S231H | -1 | 3 | 1.4 |
| | | R231K | 3 | 3 | 1.4 |
| | | D232G | 1 | 6 | 2.7 |
| | | V235A | 0 | 7 | 3.2 |
| | | V235I | 4 | 5 | 2.3 |
| | | V235T | 0 | 7 | 3.2 |
| | | E237G | 0 | 3 | 1.4 |
| | | S238A | 1 | 3 | 1.4 |
| | | I239V | 4 | 6 | 2.7 |
| | | I239L | 2 | 3 | 1.4 |
| | | Y240H | 0 | 3 | 1.4 |
| | | C242S | 0 | 3 | 1.4 |
| | | D244G | 1 | 7 | 3.2 |
| | | D244N | 2 | 3 | 1.4 |
| | | N244D | 2 | 6 | 2.7 |
| | | A246D | 0 | 3 | 1.4 |
| | | A246T | 1 | 3 | 1.4 |

| | | | | | |
|---|---|---|---|---|---|
| | | A246V | 0 | 5 | 2.3 |
| | | P247S | 1 | 3 | 1.4 |
| | | E248Q | 2 | 3 | 1.4 |
| | | R250G | -3 | 5 | 2.3 |
| | | K251R | 3 | 7 | 3.2 |
| | | Q251V | -2 | 3 | 1.4 |
| | | A252V | 0 | 8 | 3.6 |
| | | I253V | 4 | 5 | 2.3 |
| | | K254R | 3 | 9 | 4.1 |
| | | S254A | 1 | 3 | 1.4 |
| | | S255P | 1 | 4 | 1.8 |
| | | L256P | -3 | 3 | 1.4 |
| | | E258G | 0 | 4 | 1.8 |
| | | L260P | -3 | 4 | 1.8 |
| | | A272V | 0 | 3 | 1.4 |
| | | Q273P | 0 | 3 | 1.4 |
| | | T276I | 0 | 4 | 1.8 |
| | | C279R | -4 | 3 | 1.4 |
| | | S282G+ | 1 | 6 | 2.7 |
| | | S282T* | 1 | 4 | 1.8 |
| | | T287A | 1 | 3 | 1.4 |
| | | Q300R | 1 | 10 | 4.5 |
| | | Q300K | 1 | 4 | 1.8 |
| | | S300N | 1 | 10 | 4.5 |
| | | A303T | 1 | 3 | 1.4 |
| | | R304K | 3 | 3 | 1.4 |
| | | A305V | 0 | 3 | 1.4 |
| | | L308F | 2 | 3 | 1.4 |
| | | D310G | 1 | 6 | 2.7 |
| | | D310N | 2 | 6 | 2.7 |
| | | N310D | 2 | 9 | 4.1 |
| | | N310S | 1 | 4 | 1.8 |
| | | C311R | -4 | 4 | 1.8 |
| | | T312A | 1 | 6 | 2.7 |
| | | D312N | 2 | 3 | 1.4 |
| | | M313T | -1 | 3 | 1.4 |
| | | M313L | 4 | 5 | 2.3 |
| | | L314P+ | -3 | 6 | 2.7 |
| | | V315A | 0 | 3 | 1.4 |
| | | D318G | 1 | 3 | 1.4 |
| | | D319G | 1 | 3 | 1.4 |
| **0.5-0.9% of patients** | NS3 | I33V | 4 | 2 | 0.9 |
| | | V33A | 0 | 1 | 0.5 |
| | | V33T | 0 | 1 | 0.5 |
| | | Q34L | -2 | 1 | 0.5 |
| | | Q34R | 1 | 1 | 0.5 |
| | | I35V | 4 | 1 | 0.5 |
| | | V36A* | 0 | 1 | 0.5 |
| | | V36M* | 2 | 1 | 0.5 |
| | | S37P | 1 | 1 | 0.5 |
| | | T38A | 1 | 1 | 0.5 |
| | | A39S | 1 | 1 | 0.5 |
| | | V39I | 4 | 1 | 0.5 |

| | | | | |
|---|---|---|---|---|
| | A40G | 1 | 1 | 0.5 |
| | T40A | 1 | 2 | 0.9 |
| | S42A | 1 | 1 | 0.5 |
| | S42P | 1 | 1 | 0.5 |
| | A47V | 0 | 1 | 0.5 |
| | S47A | 1 | 1 | 0.5 |
| | T47I | 0 | 2 | 0.9 |
| | T47N | 0 | 1 | 0.5 |
| | C47R | -4 | 1 | 0.5 |
| | N49S | 1 | 1 | 0.5 |
| | G50S | 1 | 1 | 0.5 |
| | V51M | 2 | 1 | 0.5 |
| | C52R | -4 | 1 | 0.5 |
| | M52I | 2 | 2 | 0.9 |
| | M52V | 2 | 1 | 0.5 |
| | M52T | -1 | 1 | 0.5 |
| | T54S* | 1 | 1 | 0.5 |
| | T54A* | 1 | 2 | 0.9 |
| | V55P[+] | -1 | 1 | 0.5 |
| | Y56S[+] | -3 | 1 | 0.5 |
| | T61S | 1 | 2 | 0.9 |
| | S61P | 1 | 2 | 0.9 |
| | S61V | -1 | 1 | 0.5 |
| | K62R | 3 | 2 | 0.9 |
| | I64M | 2 | 1 | 0.5 |
| | I64T | 0 | 1 | 0.5 |
| | L64I | 2 | 2 | 0.9 |
| | A65T | 1 | 1 | 0.5 |
| | G66A | 1 | 2 | 0.9 |
| | P67Q | 0 | 1 | 0.5 |
| | S67P | 1 | 1 | 0.5 |
| | K68R | 3 | 1 | 0.5 |
| | K68S | 0 | 1 | 0.5 |
| | A71T | 1 | 2 | 0.9 |
| | I71A | -1 | 1 | 0.5 |
| | I71T | 0 | 1 | 0.5 |
| | V71I | 4 | 2 | 0.9 |
| | V71L | 2 | 1 | 0.5 |
| | L72F | 2 | 1 | 0.5 |
| | T72A | 1 | 2 | 0.9 |
| | M74T | -1 | 1 | 0.5 |
| | T76S | 1 | 1 | 0.5 |
| | V78L | 2 | 1 | 0.5 |
| | E79G | 0 | 1 | 0.5 |
| | G84S | 1 | 1 | 0.5 |
| | Q86H | 3 | 1 | 0.5 |
| | Q86L | -2 | 1 | 0.5 |
| | A87V | 0 | 1 | 0.5 |
| | A87T | 1 | 1 | 0.5 |
| | A87N | 0 | 1 | 0.5 |
| | S87G | 1 | 1 | 0.5 |
| | Q89H | 3 | 1 | 0.5 |
| | P89Q | 0 | 2 | 0.9 |

| | | | | |
|---|---|---|---|---|
| | P89A | 1 | 2 | 0.9 |
| | P89C | -3 | 2 | 0.9 |
| | P89G | -1 | 1 | 0.5 |
| | P89L | -3 | 1 | 0.5 |
| | V91G | -1 | 1 | 0.5 |
| | R92H | 2 | 1 | 0.5 |
| | R92C | -4 | 1 | 0.5 |
| | L94F | 2 | 2 | 0.9 |
| | T95I | 0 | 2 | 0.9 |
| | T95L | -2 | 1 | 0.5 |
| | T95S | 1 | 1 | 0.5 |
| | A95G | 1 | 1 | 0.5 |
| | A95T | 1 | 1 | 0.5 |
| | A95S | 1 | 2 | 0.9 |
| | P96A | 1 | 1 | 0.5 |
| | P96L | -3 | 2 | 0.9 |
| | T98G | 0 | 1 | 0.5 |
| | C99G | -3 | 1 | 0.5 |
| | C99R | -4 | 1 | 0.5 |
| | S101A | 1 | 2 | 0.9 |
| | S101G | 1 | 2 | 0.9 |
| | S101N | 1 | 1 | 0.5 |
| | A102S | 1 | 2 | 0.9 |
| | A102T | 1 | 2 | 0.9 |
| | S102L | -3 | 2 | 0.9 |
| | S102T | 1 | 1 | 0.5 |
| | D103N | 2 | 1 | 0.5 |
| | L104P | -3 | 1 | 0.5 |
| | Y105F | 7 | 2 | 0.9 |
| | Y105S | -3 | 2 | 0.9 |
| | V107A+ | 0 | 1 | 0.5 |
| | T108M | -1 | 1 | 0.5 |
| | T108P | 0 | 1 | 0.5 |
| | T108S | 1 | 1 | 0.5 |
| | H110Y | 0 | 1 | 0.5 |
| | H110N | 2 | 1 | 0.5 |
| | H110Q | 3 | 1 | 0.5 |
| | D110N | 2 | 2 | 0.9 |
| | D112N | 2 | 1 | 0.5 |
| | V113E | -2 | 1 | 0.5 |
| | P115A | 1 | 2 | 0.9 |
| | A116V | 0 | 1 | 0.5 |
| | V116G | -1 | 1 | 0.5 |
| | R117C | -4 | 2 | 0.9 |
| | R117P | 0 | 1 | 0.5 |
| | R119K | 3 | 1 | 0.5 |
| | R119Q | 1 | 1 | 0.5 |
| | G120D | 1 | 1 | 0.5 |
| | G120S | 1 | 1 | 0.5 |
| | D121G | 1 | 2 | 0.9 |
| | S122I* | -1 | 1 | 0.5 |
| | S122R* | 0 | 1 | 0.5 |
| | S122C+ | 0 | 1 | 0.5 |

| | | | | |
|---|---|---|---|---|
| | S122H+ | -1 | 1 | 0.5 |
| | T122N+ | 0 | 1 | 0.5 |
| | R123G | -3 | 1 | 0.5 |
| | T123A | 1 | 1 | 0.5 |
| | G124R | -3 | 2 | 0.9 |
| | A125S | 1 | 1 | 0.5 |
| | I127L | 2 | 2 | 0.9 |
| | L127P | -3 | 1 | 0.5 |
| | S128G | 1 | 1 | 0.5 |
| | S128P | 1 | 1 | 0.5 |
| | R130G | -3 | 2 | 0.9 |
| | I132L+ | 2 | 1 | 0.5 |
| | I132T+ | 0 | 1 | 0.5 |
| | V132L+ | 2 | 1 | 0.5 |
| | S133A | 1 | 1 | 0.5 |
| | S133P | 1 | 1 | 0.5 |
| | C134R | -4 | 1 | 0.5 |
| | C134S | 0 | 1 | 0.5 |
| | K136R | 3 | 2 | 0.9 |
| | S139P | 1 | 1 | 0.5 |
| | L143M | 4 | 1 | 0.5 |
| | V143I | 4 | 1 | 0.5 |
| | M144T | -1 | 1 | 0.5 |
| | L147M | 4 | 1 | 0.5 |
| | M147T | -1 | 1 | 0.5 |
| | S147P | 1 | 1 | 0.5 |
| | S147Q | -1 | 2 | 0.9 |
| | S147T | 1 | 1 | 0.5 |
| | A150I | -1 | 1 | 0.5 |
| | A150T | 1 | 1 | 0.5 |
| | A151V | 0 | 2 | 0.9 |
| | V151T | 0 | 1 | 0.5 |
| | I153M | 2 | 1 | 0.5 |
| | I153L | 2 | 2 | 0.9 |
| | I153T | 0 | 1 | 0.5 |
| | F154L | 2 | 2 | 0.9 |
| | F154S | -3 | 1 | 0.5 |
| | A156S* | 1 | 1 | 0.5 |
| | V158A+ | 0 | 1 | 0.5 |
| | V158E+ | -2 | 1 | 0.5 |
| | V158I* | 4 | 1 | 0.5 |
| | C159R | -4 | 1 | 0.5 |
| | V163A | 0 | 1 | 0.5 |
| | A166V | 0 | 1 | 0.5 |
| | I167V | 4 | 1 | 0.5 |
| | V167L | 2 | 2 | 0.9 |
| | V167A | 0 | 1 | 0.5 |
| | D168Y* | -4 | 2 | 0.9 |
| | D168T* | 0 | 1 | 0.5 |
| | Q168H+ | 3 | 1 | 0.5 |
| | Q168K+ | 1 | 1 | 0.5 |
| | Q168M+ | -1 | 1 | 0.5 |
| | F169L | 2 | 1 | 0.5 |

| | | | | | |
|---|---|---|---|---|---|
| | | F169S | -3 | 1 | 0.5 |
| | | I170T* | 0 | 2 | 0.9 |
| | | V170A* | 0 | 1 | 0.5 |
| | | V170M$^{+}$ | 2 | 1 | 0.5 |
| | | P171S | 1 | 1 | 0.5 |
| | | I172V | 4 | 2 | 0.9 |
| | | V172A | 0 | 2 | 0.9 |
| | | A174T | 1 | 1 | 0.5 |
| | | S174T | 1 | 1 | 0.5 |
| | | S174L | -3 | 2 | 0.9 |
| | | S174N | 1 | 1 | 0.5 |
| | | S174H | -1 | 1 | 0.5 |
| | | L175M$^{+}$ | 4 | 1 | 0.5 |
| | | E176G | 0 | 1 | 0.5 |
| | | E176A | 0 | 1 | 0.5 |
| | | S176A | 1 | 1 | 0.5 |
| | | V177I | 4 | 1 | 0.5 |
| | | T178S | 1 | 1 | 0.5 |
| | | Q178R | 1 | 2 | 0.9 |
| | | M179K | 0 | 1 | 0.5 |
| | | T179A | 1 | 1 | 0.5 |
| | | A179F | -4 | 1 | 0.5 |
| | | G24S$^{+}$ | 1 | 2 | 0.9 |
| | | K24E$^{+}$ | 0 | 1 | 0.5 |
| | | K24R* | 3 | 1 | 0.5 |
| | | K24Q$^{+}$ | 1 | 2 | 0.9 |
| | | Q24K$^{+}$ | 1 | 1 | 0.5 |
| | | Q24H* | 3 | 1 | 0.5 |
| | | S24A$^{+}$ | 1 | 1 | 0.5 |
| | | S24T$^{+}$ | 1 | 1 | 0.5 |
| | | A25S | 1 | 1 | 0.5 |
| | | K26R$^{+}$ | 3 | 2 | 0.9 |
| | | I27V | 4 | 1 | 0.5 |
| | | I27T | 0 | 1 | 0.5 |
| | | L27I | 2 | 2 | 0.9 |
| | | L27V | 2 | 1 | 0.5 |
| | | L28F* | 2 | 1 | 0.5 |
| | NS5A | L28P$^{+}$ | -3 | 1 | 0.5 |
| | | M28A* | -1 | 1 | 0.5 |
| | | M28I$^{+}$ | 2 | 2 | 0.9 |
| | | F28C$^{+}$ | -4 | 1 | 0.5 |
| | | A30L$^{+}$ | -2 | 2 | 0.9 |
| | | A30R$^{+}$ | -2 | 1 | 0.5 |
| | | L30H* | -2 | 1 | 0.5 |
| | | L30R$^{+}$ | -3 | 1 | 0.5 |
| | | R30A$^{+}$ | -2 | 1 | 0.5 |
| | | R30E$^{+}$ | -1 | 1 | 0.5 |
| | | Q30L* | -2 | 1 | 0.5 |
| | | V34F | -1 | 1 | 0.5 |
| | | L34V | 2 | 1 | 0.5 |
| | | P35S | 1 | 1 | 0.5 |
| | | V37L | 2 | 2 | 0.9 |
| | | V37M | 2 | 1 | 0.5 |

| | | | |
|---|---|---|---|
| F37I | 1 | 2 | 0.9 |
| F37Y | 7 | 1 | 0.5 |
| I37L | 2 | 1 | 0.5 |
| L37V | 2 | 2 | 0.9 |
| S38P$^{+}$ | 1 | 1 | 0.5 |
| C39R | -4 | 1 | 0.5 |
| Q40R | 1 | 1 | 0.5 |
| K41N | 1 | 2 | 0.9 |
| K41Q | 1 | 2 | 0.9 |
| K41R | 3 | 1 | 0.5 |
| Y43C | 0 | 1 | 0.5 |
| Y43F | 7 | 1 | 0.5 |
| V46E | -2 | 2 | 0.9 |
| V46M | 2 | 1 | 0.5 |
| V46T | 0 | 1 | 0.5 |
| T46V | 0 | 1 | 0.5 |
| R48L | -3 | 1 | 0.5 |
| S48P | 1 | 1 | 0.5 |
| G49A | 1 | 1 | 0.5 |
| G49E | 0 | 2 | 0.9 |
| D50E | 3 | 1 | 0.5 |
| D50G | 1 | 1 | 0.5 |
| V52G | -1 | 1 | 0.5 |
| V52I | 4 | 1 | 0.5 |
| V52L | 2 | 1 | 0.5 |
| V52M | 2 | 2 | 0.9 |
| M53V | 2 | 1 | 0.5 |
| M53I | 2 | 1 | 0.5 |
| S54H | -1 | 1 | 0.5 |
| S54L | -3 | 1 | 0.5 |
| S54T | 1 | 2 | 0.9 |
| Q54L | -2 | 1 | 0.5 |
| Q54R | 1 | 2 | 0.9 |
| Q54N | 1 | 2 | 0.9 |
| T56R | -1 | 1 | 0.5 |
| R56C | -4 | 1 | 0.5 |
| P58L$^{+}$ | -3 | 2 | 0.9 |
| P58K$^{+}$ | -1 | 1 | 0.5 |
| P58H$^{+}$ | 0 | 1 | 0.5 |
| P58Q$^{+}$ | 0 | 1 | 0.5 |
| H58D* | 1 | 1 | 0.5 |
| H58L* | -2 | 1 | 0.5 |
| H58S$^{+}$ | -1 | 1 | 0.5 |
| H58N$^{+}$ | 2 | 1 | 0.5 |
| C59W | -8 | 1 | 0.5 |
| G60S | 1 | 1 | 0.5 |
| A61V | 0 | 1 | 0.5 |
| A61S | 1 | 2 | 0.9 |
| A61T | 1 | 1 | 0.5 |
| E62D* | 3 | 2 | 0.9 |
| E62Q$^{+}$ | 2 | 1 | 0.5 |
| N62G$^{+}$ | 0 | 1 | 0.5 |
| N62S$^{+}$ | 1 | 1 | 0.5 |

| | | | | | |
|---|---|---|---|---|---|
| | | S62I+ | -1 | 2 | 0.9 |
| | | S62A+ | 1 | 2 | 0.9 |
| | | S62K+ | 0 | 1 | 0.5 |
| | | S62V+ | -1 | 2 | 0.9 |
| | | Q62D* | 2 | 1 | 0.5 |
| | | Q62R+ | 1 | 2 | 0.9 |
| | | Q62K+ | 1 | 2 | 0.9 |
| | | Q62N+ | 1 | 1 | 0.5 |
| | | I63L | 2 | 1 | 0.5 |
| | | S64P | 1 | 1 | 0.5 |
| | | H66R | 2 | 1 | 0.5 |
| | | H66Y | 0 | 1 | 0.5 |
| | | I67V | 4 | 1 | 0.5 |
| | | V67G | -1 | 1 | 0.5 |
| | | V67I | 4 | 2 | 0.9 |
| | | K68E | 0 | 2 | 0.9 |
| | | K68N | 1 | 1 | 0.5 |
| | | L69M | 4 | 1 | 0.5 |
| | | T71A | 1 | 1 | 0.5 |
| | | S71T | 1 | 1 | 0.5 |
| | | M72L | 4 | 1 | 0.5 |
| | | M72T | -1 | 2 | 0.9 |
| | | L74I | 2 | 2 | 0.9 |
| | | L74P | -3 | 1 | 0.5 |
| | | V75Y | -2 | 2 | 0.9 |
| | | A75T | 1 | 2 | 0.9 |
| | | S75A | 1 | 1 | 0.5 |
| | | T75I | 0 | 1 | 0.5 |
| | | P77A | 1 | 1 | 0.5 |
| | | P77S | 1 | 2 | 0.9 |
| | | K78N | 1 | 1 | 0.5 |
| | | T79I | 0 | 2 | 0.9 |
| | | T79M | -1 | 1 | 0.5 |
| | | T79R | -1 | 1 | 0.5 |
| | | R81K | 3 | 1 | 0.5 |
| | | R81W | 2 | 2 | 0.9 |
| | | S81R | 0 | 1 | 0.5 |
| | | M83I | 2 | 1 | 0.5 |
| | | M83L | 4 | 1 | 0.5 |
| | | W84R | 2 | 1 | 0.5 |
| | | H85L | -2 | 1 | 0.5 |
| | | H85Q | 3 | 1 | 0.5 |
| | | H85R | 2 | 1 | 0.5 |
| | | H85S | -1 | 1 | 0.5 |
| | | F88L | 2 | 1 | 0.5 |
| | | N91D | 2 | 1 | 0.5 |
| | | A92E+ | 0 | 1 | 0.5 |
| | | A92S+ | 1 | 1 | 0.5 |
| | | A92V+ | 0 | 1 | 0.5 |
| | | Y93L* | -1 | 1 | 0.5 |
| | | Y93N* | -2 | 2 | 0.9 |
| | | Y93R* | -4 | 2 | 0.9 |
| | | Y93T* | -3 | 1 | 0.5 |

| | | | | | |
|---|---|---|---|---|---|
| | | T94A | 1 | 1 | 0.5 |
| | | T94I | 0 | 1 | 0.5 |
| | | T95M | -1 | 1 | 0.5 |
| | | P97L | -3 | 1 | 0.5 |
| | | C98R | -4 | 1 | 0.5 |
| | | C98S | 0 | 2 | 0.9 |
| | | T99A | 1 | 2 | 0.9 |
| | | T99V | 0 | 2 | 0.9 |
| | | V99A | 0 | 2 | 0.9 |
| | | V99M | 2 | 1 | 0.5 |
| | | P100A | 1 | 2 | 0.9 |
| | | I101A | -1 | 1 | 0.5 |
| | | I101T | 0 | 2 | 0.9 |
| | | C101Y | 0 | 1 | 0.5 |
| | | S101L | -3 | 1 | 0.5 |
| | | S101A | 1 | 1 | 0.5 |
| | | S101T | 1 | 1 | 0.5 |
| | | P102S | 1 | 1 | 0.5 |
| | | A103T | 1 | 1 | 0.5 |
| | | T103A | 1 | 1 | 0.5 |
| | | P104C | -3 | 1 | 0.5 |
| | | P104S | 1 | 1 | 0.5 |
| | | P104L | -3 | 1 | 0.5 |
| | | P104H | 0 | 2 | 0.9 |
| | | N105H | 2 | 1 | 0.5 |
| | | T107E | 0 | 1 | 0.5 |
| | | T107M | -1 | 1 | 0.5 |
| | | T107N | 0 | 1 | 0.5 |
| | | T107S | 1 | 2 | 0.9 |
| | | S107V | -1 | 1 | 0.5 |
| | | S107E | 0 | 1 | 0.5 |
| | | S107F | -3 | 1 | 0.5 |
| | | S107G | 1 | 1 | 0.5 |
| | | S107P | 1 | 2 | 0.9 |
| | | R108G | -3 | 1 | 0.5 |
| | | R108F | -4 | 1 | 0.5 |
| | | F108S | -3 | 1 | 0.5 |
| | | F108V | -1 | 1 | 0.5 |
| | | L110P | -3 | 1 | 0.5 |
| | | L110I | 2 | 1 | 0.5 |
| | | L110M | 4 | 1 | 0.5 |
| | | W111L | -2 | 2 | 0.9 |
| | | W111- | - | 1 | 0.5 |
| | | R112C | -4 | 1 | 0.5 |
| | | R112G | -3 | 2 | 0.9 |
| | | V113L | 2 | 1 | 0.5 |
| | | S114T | 1 | 2 | 0.9 |
| | | S116T | 1 | 1 | 0.5 |
| | | N116T | 0 | 1 | 0.5 |
| | | E117G | 0 | 1 | 0.5 |
| | | V119I | 4 | 1 | 0.5 |
| | | E120G | 0 | 1 | 0.5 |
| | | R122K | 3 | 2 | 0.9 |

| | | | | | |
|---|---|---|---|---|---|
| | | T122K | 0 | 1 | 0.5 |
| | | R123K | 3 | 1 | 0.5 |
| | | V124A | 0 | 1 | 0.5 |
| | | V124E | -2 | 1 | 0.5 |
| | | V124K | -2 | 1 | 0.5 |
| | | V124L | 2 | 1 | 0.5 |
| | | V124M | 2 | 1 | 0.5 |
| | | V124I | 4 | 1 | 0.5 |
| | | G125S | 1 | 1 | 0.5 |
| | | E126G | 0 | 1 | 0.5 |
| | | P126S | 1 | 1 | 0.5 |
| | | F127H | -2 | 1 | 0.5 |
| | | F127A | -4 | 1 | 0.5 |
| | | F127C | -4 | 1 | 0.5 |
| | | F127Y | 7 | 1 | 0.5 |
| | | Y129F | 7 | 1 | 0.5 |
| | | V130I | 4 | 2 | 0.9 |
| | | G132A | 1 | 2 | 0.9 |
| | | M133L | 4 | 1 | 0.5 |
| | | T134A | 1 | 1 | 0.5 |
| | | T135I | 0 | 1 | 0.5 |
| | | T135S | 1 | 2 | 0.9 |
| | | D136N | 2 | 1 | 0.5 |
| | | E137A | 0 | 1 | 0.5 |
| | | E137D | 3 | 1 | 0.5 |
| | | N137S | 1 | 1 | 0.5 |
| | | L138P | -3 | 1 | 0.5 |
| | | C140G | -3 | 2 | 0.9 |
| | | V140L | 2 | 1 | 0.5 |
| | | Q143R | 1 | 1 | 0.5 |
| | | V144I | 4 | 1 | 0.5 |
| | | P145L | -3 | 1 | 0.5 |
| | | P145S | 1 | 1 | 0.5 |
| | | S146A | 1 | 1 | 0.5 |
| | | S146P | 1 | 2 | 0.9 |
| | | A146G | 1 | 2 | 0.9 |
| | | E148D | 3 | 1 | 0.5 |
| | | F149S | -3 | 1 | 0.5 |
| | | F150L | 2 | 1 | 0.5 |
| | | F150S | -3 | 1 | 0.5 |
| | | T151A | 1 | 1 | 0.5 |
| | NS5B | K124T | 0 | 1 | 0.5 |
| | | K124N | 1 | 1 | 0.5 |
| | | K124- | - | 1 | 0.5 |
| | | E124R | -1 | 1 | 0.5 |
| | | L126M | 4 | 1 | 0.5 |
| | | L126S | -3 | 1 | 0.5 |
| | | L127P | -3 | 1 | 0.5 |
| | | L127Q | -2 | 1 | 0.5 |
| | | R127L | -3 | 1 | 0.5 |
| | | E128V | -2 | 1 | 0.5 |
| | | E128G | 0 | 2 | 0.9 |
| | | D129N | 2 | 2 | 0.9 |

| | | | |
|---|---|---|---|
| N130K | 1 | 1 | 0.5 |
| S130C | 0 | 1 | 0.5 |
| T130A | 1 | 2 | 0.9 |
| T130I | 0 | 1 | 0.5 |
| V131I | 4 | 1 | 0.5 |
| E131R | -1 | 1 | 0.5 |
| E131Q | 2 | 2 | 0.9 |
| E131A | 0 | 1 | 0.5 |
| E131G | 0 | 1 | 0.5 |
| T131A | 1 | 1 | 0.5 |
| T131N | 0 | 1 | 0.5 |
| T131V | 0 | 1 | 0.5 |
| I134T | 0 | 2 | 0.9 |
| I134V | 4 | 1 | 0.5 |
| D135N | 2 | 1 | 0.5 |
| D135Q | 2 | 1 | 0.5 |
| P135S | 1 | 2 | 0.9 |
| Q135E | 2 | 1 | 0.5 |
| T136S | 1 | 1 | 0.5 |
| T136A | 1 | 1 | 0.5 |
| T137A | 1 | 1 | 0.5 |
| M139V | 2 | 2 | 0.9 |
| N142D | 2 | 1 | 0.5 |
| N142T | 0 | 2 | 0.9 |
| E143D | 3 | 1 | 0.5 |
| E143Q | 2 | 1 | 0.5 |
| V144A | 0 | 1 | 0.5 |
| F145Y | 7 | 1 | 0.5 |
| C146Y | 0 | 1 | 0.5 |
| V147L | 2 | 2 | 0.9 |
| V147M | 2 | 2 | 0.9 |
| Q148- | - | 1 | 0.5 |
| Q148P | 0 | 1 | 0.5 |
| D148E | 3 | 1 | 0.5 |
| D148S | 0 | 1 | 0.5 |
| E150D | 3 | 1 | 0.5 |
| E150G | 0 | 1 | 0.5 |
| E150K | 0 | 1 | 0.5 |
| V150S | -1 | 2 | 0.9 |
| K151E | 0 | 2 | 0.9 |
| K151T | 0 | 1 | 0.5 |
| G153S | 1 | 1 | 0.5 |
| R154G | -3 | 1 | 0.5 |
| R154H | 2 | 1 | 0.5 |
| K155E | 0 | 2 | 0.9 |
| K155R | 3 | 1 | 0.5 |
| P156S | 1 | 1 | 0.5 |
| P156A | 1 | 2 | 0.9 |
| A157V | 0 | 1 | 0.5 |
| R158H | 2 | 1 | 0.5 |
| I160T | 0 | 1 | 0.5 |
| I160V | 4 | 2 | 0.9 |
| V161A | 0 | 2 | 0.9 |

| | | | | |
|---|---|---|---|---|
| | Y162F | 7 | 1 | 0.5 |
| | F162S | -3 | 1 | 0.5 |
| | D164G | 1 | 2 | 0.9 |
| | G166S | 1 | 1 | 0.5 |
| | V167I | 4 | 1 | 0.5 |
| | V167M | 2 | 1 | 0.5 |
| | R168H | 2 | 1 | 0.5 |
| | V169G | -1 | 1 | 0.5 |
| | V169L | 2 | 1 | 0.5 |
| | V169A | 0 | 1 | 0.5 |
| | E171K | 0 | 1 | 0.5 |
| | K172R | 3 | 1 | 0.5 |
| | R173K | 3 | 1 | 0.5 |
| | M173I | 2 | 1 | 0.5 |
| | M173R | 0 | 1 | 0.5 |
| | M173T | -1 | 1 | 0.5 |
| | A174T | 1 | 1 | 0.5 |
| | L175F | 2 | 1 | 0.5 |
| | L175M | 4 | 1 | 0.5 |
| | Y176C | 0 | 1 | 0.5 |
| | Y176H | 0 | 1 | 0.5 |
| | D177E | 3 | 1 | 0.5 |
| | D177S | 0 | 1 | 0.5 |
| | A178V | 0 | 1 | 0.5 |
| | V178Q | -2 | 1 | 0.5 |
| | V178I | 4 | 1 | 0.5 |
| | V178L | 2 | 2 | 0.9 |
| | A179T | 1 | 1 | 0.5 |
| | I179L | 2 | 1 | 0.5 |
| | I179V | 4 | 1 | 0.5 |
| | V179A | 0 | 2 | 0.9 |
| | T179A | 1 | 1 | 0.5 |
| | S180G | 1 | 2 | 0.9 |
| | S180L | -3 | 1 | 0.5 |
| | S180R | 0 | 1 | 0.5 |
| | S180T | 1 | 2 | 0.9 |
| | K180E | 0 | 1 | 0.5 |
| | K180N | 1 | 1 | 0.5 |
| | Q180E | 2 | 1 | 0.5 |
| | K181Q | 1 | 1 | 0.5 |
| | K181R | 3 | 1 | 0.5 |
| | Q181K | 1 | 2 | 0.9 |
| | T181I | 0 | 1 | 0.5 |
| | L182P | -3 | 2 | 0.9 |
| | S183P | 1 | 1 | 0.5 |
| | P183S | 1 | 1 | 0.5 |
| | E184K | 0 | 1 | 0.5 |
| | L184M | 4 | 1 | 0.5 |
| | L184T | -2 | 1 | 0.5 |
| | I184N | -2 | 1 | 0.5 |
| | I184V | 4 | 1 | 0.5 |
| | Q184R | 1 | 1 | 0.5 |
| | Q184H | 3 | 2 | 0.9 |

| | | | | |
|---|---|---|---|---|
| | Q184I | -2 | 1 | 0.5 |
| | Q184L | -2 | 1 | 0.5 |
| | Q184V | -2 | 2 | 0.9 |
| | A185V | 0 | 2 | 0.9 |
| | A185T | 1 | 1 | 0.5 |
| | T186A | 1 | 1 | 0.5 |
| | V186A | 0 | 2 | 0.9 |
| | V186G | -1 | 1 | 0.5 |
| | V186L | 2 | 1 | 0.5 |
| | M187I | 2 | 2 | 0.9 |
| | M187T | -1 | 1 | 0.5 |
| | M187V | 2 | 1 | 0.5 |
| | G188D | 1 | 2 | 0.9 |
| | G188R | -3 | 1 | 0.5 |
| | D189G | 1 | 2 | 0.9 |
| | E189K | 0 | 1 | 0.5 |
| | S189F | -3 | 1 | 0.5 |
| | S189G | 1 | 1 | 0.5 |
| | S189R | 0 | 1 | 0.5 |
| | A189E | 0 | 1 | 0.5 |
| | A189T | 1 | 1 | 0.5 |
| | A190S | 1 | 2 | 0.9 |
| | S190P | 1 | 1 | 0.5 |
| | S190T | 1 | 2 | 0.9 |
| | Y191C | 0 | 1 | 0.5 |
| | G192R | -3 | 1 | 0.5 |
| | F193L | 2 | 1 | 0.5 |
| | F193S | -3 | 2 | 0.9 |
| | Q194R | 1 | 2 | 0.9 |
| | Y195C | 0 | 2 | 0.9 |
| | S196L | -3 | 1 | 0.5 |
| | P197S | 1 | 1 | 0.5 |
| | G198R | -3 | 1 | 0.5 |
| | G198K | -2 | 2 | 0.9 |
| | G198Q | -1 | 1 | 0.5 |
| | S198A | 1 | 1 | 0.5 |
| | E202K | 0 | 1 | 0.5 |
| | F203L | 2 | 2 | 0.9 |
| | F203S | -3 | 1 | 0.5 |
| | R203L | -3 | 1 | 0.5 |
| | L204P | -3 | 1 | 0.5 |
| | L205F | 2 | 1 | 0.5 |
| | L205M | 4 | 1 | 0.5 |
| | Q206H | 3 | 1 | 0.5 |
| | Q206K | 1 | 1 | 0.5 |
| | Q206L | -2 | 1 | 0.5 |
| | K206R | 3 | 2 | 0.9 |
| | K206G | -2 | 1 | 0.5 |
| | N206T | 0 | 2 | 0.9 |
| | N206G | 0 | 1 | 0.5 |
| | E206K | 0 | 1 | 0.5 |
| | E206R | -1 | 1 | 0.5 |
| | A207G | 1 | 1 | 0.5 |

| | | | | |
|---|---|---|---|---|
| | M207T | -1 | 2 | 0.9 |
| | W208R | 2 | 2 | 0.9 |
| | K209S | 0 | 2 | 0.9 |
| | K209E | 0 | 2 | 0.9 |
| | K209T | 0 | 1 | 0.5 |
| | T209A | 1 | 2 | 0.9 |
| | S210F | -3 | 1 | 0.5 |
| | S210G | 1 | 1 | 0.5 |
| | S210H | -1 | 2 | 0.9 |
| | S210L | -3 | 1 | 0.5 |
| | S210P | 1 | 1 | 0.5 |
| | S210R | 0 | 2 | 0.9 |
| | S210A | 1 | 2 | 0.9 |
| | S210T | 1 | 1 | 0.5 |
| | K212N | 1 | 1 | 0.5 |
| | T213I | 0 | 2 | 0.9 |
| | T213N | 0 | 2 | 0.9 |
| | T213S | 1 | 1 | 0.5 |
| | T213V | 0 | 1 | 0.5 |
| | T213A | 1 | 2 | 0.9 |
| | S213R | 0 | 2 | 0.9 |
| | L215S | -3 | 1 | 0.5 |
| | M215L | 4 | 2 | 0.9 |
| | M215T | -1 | 1 | 0.5 |
| | G216A | 1 | 1 | 0.5 |
| | F217L | 2 | 2 | 0.9 |
| | F217S | -3 | 1 | 0.5 |
| | S218A | 1 | 1 | 0.5 |
| | S218P | 1 | 2 | 0.9 |
| | Y219C | 0 | 1 | 0.5 |
| | Y219H | 0 | 1 | 0.5 |
| | D220G | 1 | 2 | 0.9 |
| | D220V | -2 | 1 | 0.5 |
| | T221A | 1 | 1 | 0.5 |
| | R222C | -4 | 1 | 0.5 |
| | R222H | 2 | 1 | 0.5 |
| | C223R | -4 | 2 | 0.9 |
| | D225G | 1 | 2 | 0.9 |
| | T229A | 1 | 1 | 0.5 |
| | E230K | 0 | 2 | 0.9 |
| | Q231R | 1 | 2 | 0.9 |
| | S231R | 0 | 1 | 0.5 |
| | R231G | -3 | 1 | 0.5 |
| | R231S | 0 | 2 | 0.9 |
| | I233T | 0 | 1 | 0.5 |
| | I233V | 4 | 1 | 0.5 |
| | T235V | 0 | 1 | 0.5 |
| | T235A | 1 | 1 | 0.5 |
| | A235T | 1 | 1 | 0.5 |
| | V235G | -1 | 1 | 0.5 |
| | V235M | 2 | 1 | 0.5 |
| | E236K | 0 | 1 | 0.5 |
| | E236G | 0 | 1 | 0.5 |

| | | | | |
|---|---|---|---|---|
| | E236V | -2 | 1 | 0.5 |
| | A238V | 0 | 2 | 0.9 |
| | S238P | 1 | 2 | 0.9 |
| | I239T | 0 | 2 | 0.9 |
| | Y240C | 0 | 1 | 0.5 |
| | Q241R | 1 | 2 | 0.9 |
| | C242Y | 0 | 1 | 0.5 |
| | N244S | 1 | 2 | 0.9 |
| | N244T | 0 | 1 | 0.5 |
| | L245M | 4 | 2 | 0.9 |
| | L245S | -3 | 1 | 0.5 |
| | L245P | -3 | 2 | 0.9 |
| | L245V | 2 | 1 | 0.5 |
| | D246A | 0 | 1 | 0.5 |
| | E246D | 3 | 2 | 0.9 |
| | A246I | -1 | 1 | 0.5 |
| | P247L | -3 | 2 | 0.9 |
| | Q248R | 1 | 1 | 0.5 |
| | E248D | 3 | 2 | 0.9 |
| | A249P | 1 | 1 | 0.5 |
| | A249V | 0 | 1 | 0.5 |
| | R250M | 0 | 1 | 0.5 |
| | V251A | 0 | 2 | 0.9 |
| | V251M | 2 | 1 | 0.5 |
| | K251E | 0 | 2 | 0.9 |
| | Q251R | 1 | 2 | 0.9 |
| | A252T | 1 | 1 | 0.5 |
| | V252M | 2 | 1 | 0.5 |
| | V252A | 0 | 1 | 0.5 |
| | I253M | 2 | 1 | 0.5 |
| | I253T | 0 | 2 | 0.9 |
| | K254N | 1 | 1 | 0.5 |
| | R254G | -3 | 2 | 0.9 |
| | T254A | 1 | 1 | 0.5 |
| | S254P | 1 | 1 | 0.5 |
| | S254T | 1 | 1 | 0.5 |
| | S255F | -3 | 1 | 0.5 |
| | S255A | 1 | 2 | 0.9 |
| | T257S | 1 | 1 | 0.5 |
| | T257A | 1 | 1 | 0.5 |
| | E258A | 0 | 1 | 0.5 |
| | E258Q | 2 | 1 | 0.5 |
| | E258K | 0 | 2 | 0.9 |
| | R259G | -3 | 1 | 0.5 |
| | R259L | -3 | 1 | 0.5 |
| | Y261H | 0 | 2 | 0.9 |
| | V262I | 4 | 2 | 0.9 |
| | C262R | -4 | 1 | 0.5 |
| | I262T | 0 | 2 | 0.9 |
| | G263S | 1 | 1 | 0.5 |
| | P265S | 1 | 1 | 0.5 |
| | L266M | 4 | 1 | 0.5 |
| | L266P | -3 | 1 | 0.5 |

| | | | | |
|---|---|---|---|---|
| | T267A | 1 | 2 | 0.9 |
| | T267I | 0 | 2 | 0.9 |
| | Y267C | 0 | 1 | 0.5 |
| | Y267F | 7 | 1 | 0.5 |
| | F267L | 2 | 2 | 0.9 |
| | N268D | 2 | 1 | 0.5 |
| | K270E | 0 | 2 | 0.9 |
| | K270R | 3 | 2 | 0.9 |
| | K270Q | 1 | 1 | 0.5 |
| | G271E | 0 | 1 | 0.5 |
| | G271R | -3 | 1 | 0.5 |
| | D272G | 1 | 1 | 0.5 |
| | E272G | 0 | 2 | 0.9 |
| | A272T | 1 | 1 | 0.5 |
| | A272D | 0 | 1 | 0.5 |
| | Q272E | 2 | 1 | 0.5 |
| | Q272R | 1 | 1 | 0.5 |
| | N273H | 2 | 1 | 0.5 |
| | N273T | 0 | 2 | 0.9 |
| | N273D | 2 | 2 | 0.9 |
| | T276V | 0 | 2 | 0.9 |
| | Y276C | 0 | 1 | 0.5 |
| | Y276H | 0 | 2 | 0.9 |
| | R277C | -4 | 1 | 0.5 |
| | R278G | -3 | 1 | 0.5 |
| | R278K | 3 | 1 | 0.5 |
| | R278W | 2 | 1 | 0.5 |
| | C279Y | 0 | 1 | 0.5 |
| | G283S | 1 | 1 | 0.5 |
| | L285P | -3 | 2 | 0.9 |
| | L285F | 2 | 1 | 0.5 |
| | F285L | 2 | 1 | 0.5 |
| | F285Y | 7 | 1 | 0.5 |
| | F285- | - | 1 | 0.5 |
| | P286L | -3 | 1 | 0.5 |
| | T286A | 1 | 1 | 0.5 |
| | T286M | -1 | 1 | 0.5 |
| | S288G | 1 | 2 | 0.9 |
| | S288I | -1 | 1 | 0.5 |
| | C289R[+] | -4 | 1 | 0.5 |
| | F289L* | 2 | 1 | 0.5 |
| | G290D | 1 | 1 | 0.5 |
| | N291D | 2 | 1 | 0.5 |
| | N291S | 1 | 2 | 0.9 |
| | I293L | 2 | 1 | 0.5 |
| | I293V | 4 | 1 | 0.5 |
| | L293M | 4 | 1 | 0.5 |
| | T294A | 1 | 2 | 0.9 |
| | C295R | -4 | 2 | 0.9 |
| | Y296C | 0 | 1 | 0.5 |
| | I297V | 4 | 1 | 0.5 |
| | L297I | 2 | 1 | 0.5 |
| | L297P | -3 | 1 | 0.5 |

| | | | | | |
|---|---|---|---|---|---|
| | | L297S | -3 | 2 | 0.9 |
| | | K298R | 3 | 2 | 0.9 |
| | | A299T | 1 | 1 | 0.5 |
| | | Q300F | -5 | 1 | 0.5 |
| | | Q300L | -2 | 1 | 0.5 |
| | | Q300N | 1 | 1 | 0.5 |
| | | S300V | -1 | 1 | 0.5 |
| | | S300Q | -1 | 1 | 0.5 |
| | | R300K | 3 | 1 | 0.5 |
| | | T300I | 0 | 1 | 0.5 |
| | | T300M | -1 | 1 | 0.5 |
| | | A301V | 0 | 1 | 0.5 |
| | | A302S | 1 | 2 | 0.9 |
| | | C303Y | 0 | 1 | 0.5 |
| | | C303R | -4 | 1 | 0.5 |
| | | I303T | 0 | 2 | 0.9 |
| | | R304- | - | 1 | 0.5 |
| | | K304E | 0 | 1 | 0.5 |
| | | A305T | 1 | 2 | 0.9 |
| | | A306T | 1 | 2 | 0.9 |
| | | A306V | 0 | 1 | 0.5 |
| | | G307D | 1 | 1 | 0.5 |
| | | G307R | -3 | 2 | 0.9 |
| | | G307A | 1 | 1 | 0.5 |
| | | G307K | -2 | 1 | 0.5 |
| | | G307S | 1 | 2 | 0.9 |
| | | K307G | -2 | 1 | 0.5 |
| | | K307E | 0 | 2 | 0.9 |
| | | L308P | -3 | 2 | 0.9 |
| | | K309R | 3 | 1 | 0.5 |
| | | Q309W | -5 | 2 | 0.9 |
| | | R309Q | 1 | 1 | 0.5 |
| | | V309I | 4 | 1 | 0.5 |
| | | D310E | 3 | 1 | 0.5 |
| | | D310A | 0 | 1 | 0.5 |
| | | N310A | 0 | 2 | 0.9 |
| | | C311Y | 0 | 2 | 0.9 |
| | | C311H | -3 | 1 | 0.5 |
| | | C311S | 0 | 1 | 0.5 |
| | | P311A | 1 | 1 | 0.5 |
| | | T312I | 0 | 2 | 0.9 |
| | | T312S | 1 | 1 | 0.5 |
| | | T312M | -1 | 1 | 0.5 |
| | | D312E | 3 | 1 | 0.5 |
| | | F313L | 2 | 1 | 0.5 |
| | | M313I | 2 | 1 | 0.5 |
| | | M313S | -2 | 1 | 0.5 |
| | | M313V | 2 | 2 | 0.9 |
| | | L314F+ | 2 | 1 | 0.5 |
| | | L314I+ | 2 | 1 | 0.5 |
| | | C316Y* | 0 | 2 | 0.9 |
| | | C316R+ | -4 | 2 | 0.9 |
| | | C316S+ | 0 | 1 | 0.5 |

| | | | | | |
|---|---|---|---|---|---|
| | | G317S | 1 | 1 | 0.5 |
| | | D318V | -2 | 1 | 0.5 |
| | | L320P[+] | -3 | 1 | 0.5 |

[a] The clinical history of the 220 patients cohort under study was described in Chen et al., Antiviral Research (2020) 174: 10469.

[b] The HCV genome residue numbering corresponds to the H77 genome (accession number #AF009606). The reference sequences for each substitution was described in Soria et al., BMC infectious and diseases (2018) 18(1):446.

[c] PAM250 substitution matrix values are taken from Feng and Doolittle (Methods in Enzymology 266: 368-382, 1996): PAM250<0, lower acceptability than expected, meaning a rare replacement; PAM250=0, acceptability as expected; PAM250>0 acceptability higher than expected.

* RAS, resistance-associated substitutions

[+] RAP, putative new resistance-associated substitutions, previously described in Chen et al., Antiviral Research (2020) 174: 104694. In this paper, for calculations RAP are considered as RAS for simplicity.

- A minus symbol after the number means stop codon.

**Table S2. Statistical analysis between the HRS combinations observed in patients carrying at least one HRS.**

| G1a | | | |
|---|---|---|---|
| **Combination 1[a]** | **Combination 2** | **p-value** | **Significance[b]** |
| **R78K + Q309R** | R78K | 0.008397 | ** |
| | S231N | 4.385 x $10^{-5}$ | *** |
| | Q309R | 0.0001677 | *** |
| | T64A + R78K | 0.0001677 | *** |
| | T64A + S231N | 4.385 x $10^{-5}$ | *** |
| | R78K + S231N | 0.0005434 | *** |
| | T64A + R78K + S231N | 4.385 x $10^{-5}$ | *** |
| | T64A + R78K + Q309R | 0.2544 | **ns** |
| | R78K + S231N + Q309R | 0.0001677 | *** |
| **T64A + R78K + Q309R** | R78K | 0.06215 | **ns** |
| | S231N | 0.000734 | *** |
| | Q309R | 0.00247 | ** |
| | T64A + R78K | 0.00247 | ** |
| | T64A + S231N | 0.000734 | *** |
| | R78K + S231N | 0.006855 | ** |
| | R78K + Q309R | 0.7456 | **ns** |
| | T64A + R78K + S231N | 0.000734 | *** |
| | R78K + S231N + Q309R | 0.00247 | ** |
| **G1b** | | | |
| **Combination 1[a]** | **Combination 2** | **p-value** | **Significance[b]** |
| **S213C + A218S** | S213C | 0.000178 | *** |
| | A218S | 0.003564 | ** |
| | S231N | 4.915 x $10^{-5}$ | *** |
| | T64A + S213C | 4.915 x $10^{-5}$ | *** |
| | T64A + A218S | 0.0005507 | *** |
| | A218S + S231N | 0.000178 | *** |
| | A218S + Q309R | 4.915 x $10^{-5}$ | *** |
| | S213C + S231N | 0.003564 | ** |
| | S213C + Q309R | 4.915 x $10^{-5}$ | *** |
| | T64A + S213C + A218S | 0.0005507 | *** |
| | T64A + S213C + S231N | 0.000178 | *** |
| | T64A + A218S + S231N | 0.000178 | *** |
| | T64A + S231N + Q309R | 4.915 x $10^{-5}$ | *** |
| | S213C + A218S + S231N | 0.5 | **ns** |
| | S213C + A218S + Q309R | 0.0005507 | *** |
| | T64A + S213C + A218S + S231N | 0.1106 | **ns** |
| | T64A + S213C + A218S + Q309R | 4.915 x $10^{-5}$ | *** |
| | S213C + A218S + S231N + Q309R | 0.000178 | *** |
| | T64A + S213C + A218S + S231N + Q309R | 0.0005507 | *** |
| **S213C + A218S + S231N** | S213C | 0.000178 | *** |
| | A218S | 0.003564 | ** |
| | S231N | 4.915 x $10^{-5}$ | *** |
| | T64A + S213C | 4.915 x $10^{-5}$ | *** |

| | | | |
|---|---|---|---|
| | T64A + S231N | 0.0005507 | *** |
| | A218S + S231N | 0.000178 | *** |
| | A218S + Q309R | 4.915 x $10^{-5}$ | *** |
| | S213C + A218S | 0.5 | **ns** |
| | S213C + S231N | 0.003564 | ** |
| | S213C + Q309R | 4.915 x $10^{-5}$ | *** |
| | T64A + S213C + A218S | 0.0005507 | *** |
| | T64A + S213C + S231N | 0.000178 | *** |
| | T64A + A218S + S231N | 0.000178 | *** |
| | T64A + S231N + Q309R | 4.915 x $10^{-5}$ | *** |
| | S213C + A218S + Q309R | 0.0005507 | *** |
| | T64A + S213C + A218S + S231N | 0.1106 | **ns** |
| | T64A + S213C + A218S + Q309R | 4.915 x $10^{-5}$ | *** |
| | S213C + A218S + S231N + Q309R | 0.000178 | *** |
| | T64A + S213C + A218S + S231N + Q309R | 0.0005507 | *** |
| **T64A + S213C + A218S + S231N** | S213C | 0.01048 | * |
| | A218S | 0.09454 | **ns** |
| | S231N | 0.00352 | ** |
| | T64A + S213C | 0.00352 | ** |
| | T64A + S231N | 0.02541 | * |
| | A218S + S231N | 0.01048 | * |
| | A218S + Q309R | 0.00352 | ** |
| | S213C + A218S | 0.8894 | **ns** |
| | S213C + S231N | 0.09454 | **ns** |
| | S213C + Q309R | 0.00352 | ** |
| | T64A + S213C + A218S | 0.02541 | * |
| | T64A + S213C + S231N | 0.01048 | * |
| | T64A + A218S + S231N | 0.01048 | * |
| | T64A + S231N + Q309R | 0.00352 | ** |
| | S213C + A218S + S231N | 0.8894 | **ns** |
| | S213C + A218S + Q309R | 0.02541 | * |
| | T64A + S213C + A218S + Q309R | 0.00352 | ** |
| | S213C + A218S + S231N + Q309R | 0.01048 | * |
| | T64A + S213C + A218S + S231N + Q309R | 0.02541 | * |

[a] Combinations 1 for G1a and G1b represent those HRS combinations with the higher percentage of patients.

[b] The statistical significance of the differences is given as follows: ns=not significant; * $p<0.05$; ** $p<0.01$; *** $p<0.001$.

**Table S3. Statistical analysis for the difference between the frequency of individual HRS in HCV from patients in the basal (pre-treatment) and post-treatment (failure) samples.**

| G1a | Comparison | p-value | Significance[a] |
|---|---|---|---|
| T64A | Post-treatment = pre-treatment<br>vs<br>Post-treatment > pre-treatment | na | na |
| | Post-treatment = pre-treatment<br>vs<br>Post-treatment < pre-treatment | na | na |
| R78K | Post-treatment = pre-treatment<br>vs<br>Post-treatment > pre-treatment | 0.001362 | ** |
| | Post-treatment = pre-treatment<br>vs<br>Post-treatment < pre-treatment | 0.001362 | ** |
| S231N | Post-treatment = pre-treatment<br>vs<br>Post-treatment > pre-treatment | na | na |
| | Post-treatment = pre-treatment<br>vs<br>Post-treatment < pre-treatment | na | na |
| Q309R | Post-treatment = pre-treatment<br>vs<br>Post-treatment > pre-treatment | 0.00186 | ** |
| | Post-treatment = pre-treatment<br>vs<br>Post-treatment < pre-treatment | 0.02967 | * |
| **G1b** | **Comparison** | **p-value** | **Significance** |
| T64A | Post-treatment = pre-treatment<br>vs<br>Post-treatment > pre-treatment | 0.06483 | ns |
| | Post-treatment = pre-treatment<br>vs<br>Post-treatment < pre-treatment | 0.1318 | ns |
| S213C | Post-treatment = pre-treatment<br>vs<br>Post-treatment > pre-treatment | $3.405 \times 10^{-12}$ | *** |
| | Post-treatment = pre-treatment<br>vs<br>Post-treatment < pre-treatment | $3.405 \times 10^{-12}$ | *** |
| A218S | Post-treatment = pre-treatment<br>vs<br>Post-treatment > pre-treatment | $1.992 \times 10^{-9}$ | *** |

| | | | |
|---|---|---|---|
| | Post-treatment = pre-treatment<br>vs<br>Post-treatment < pre-treatment | $1.68 \times 10^{-7}$ | *** |
| S231N | Post-treatment = pre-treatment<br>vs<br>Post-treatment > pre-treatment | $2.556 \times 10^{-5}$ | *** |
| | Post-treatment = pre-treatment<br>vs<br>Post-treatment < pre-treatment | $2.556 \times 10^{-5}$ | *** |
| Q309R | Post-treatment = pre-treatment<br>vs<br>Post-treatment > pre-treatment | 0.1567 | ns |
| | Post-treatment = pre-treatment<br>vs<br>Post-treatment < pre-treatment | 0.06067 | ns |

[a] The statistical significance of the differences is given as follows: na=not applicable; ns=not significant; * p<0.05; ** p<0.01; *** p<0.001.

**Table S4. Statistical analysis of the association between RAS L159F and C316N (in bold) with HRS.**

| Individual HRS | | | |
|---|---|---|---|
| **Combination 1** | **Combination 2** | **p-value** | **Significance**[a] |
| | **C316N** + T64A | $4.479 \times 10^{-14}$ | *** |
| | **C316N** + R78K | $< 2.2 \times 10^{-16}$ | *** |
| **C316N** + A218S | **C316N** + S213C | $1.785 \times 10^{-2}$ | * |
| | **C316N** + S231N | $1.165 \times 10^{-7}$ | *** |
| | **C316N** + Q309R | $< 2.2 \times 10^{-16}$ | *** |
| | **C316N** + T64A | $6.357 \times 10^{-10}$ | *** |
| | **C316N** + R78K | $< 2.2 \times 10^{-16}$ | *** |
| **C316N** + S213C | **C316N** + A218S | 0.9822 | **ns** |
| | **C316N** + S231N | $2.565 \times 10^{-4}$ | *** |
| | **C316N** + Q309R | $4.657 \times 10^{-13}$ | *** |
| | **L159F** + T64A | $5.089 \times 10^{-12}$ | *** |
| | **L159F** + R78K | $< 2.2 \times 10^{-16}$ | *** |
| **L159F** + A218S | **L159F** + S213C | 0.1335 | **ns** |
| | **L159F** + S231N | $7.489 \times 10^{-5}$ | *** |
| | **L159F** + Q309R | $4.803 \times 10^{-15}$ | *** |
| | **L159F** + T64A | $2.137 \times 10^{-9}$ | *** |
| | **L159F** + R78K | $7.884 \times 10^{-16}$ | *** |
| **L159F** + S213C | **L159F** + A218S | 0.8665 | **ns** |
| | **L159F** + S231N | $4.504 \times 10^{-3}$ | ** |
| | **L159F** + Q309R | $2.972 \times 10^{-12}$ | *** |

| HRS combinations | | | |
|---|---|---|---|
| **Combination 1** | **Combination 2** | **p-value** | **Significance** |
| | **C316N** + R78K + Q309R | $9.799 \times 10^{-5}$ | *** |
| **C316N** + S213C + | **C316N** + T64A + R78K + Q309R | $9.799 \times 10^{-5}$ | *** |
| A218S | **C316N** + S213C + A218S + S231N | 0.5 | **ns** |
| | **C316N** + T64A + S213C + A218S + S231N | 0.1736 | **ns** |
| | **C316N** + R78K + Q309R | $9.799 \times 10^{-5}$ | *** |
| **C316N** + S213C + | **C316N** + T64A + R78K + Q309R | $9.799 \times 10^{-5}$ | *** |
| A218S + S231N | **C316N** + S213C + A218S | 0.5 | **ns** |
| | **C316N** + T64A + S213C + A218S + S231N | 0.1736 | **ns** |
| | **L159F** + R78K + Q309R | $3.305 \times 10^{-4}$ | *** |
| **L159F** + S213C + | **L159F** + T64A + R78K + Q309R | $3.305 \times 10^{-4}$ | *** |
| A218S | **L159F** + S213C + A218S + S231N | 0.5 | **ns** |
| | **L159F** + T64A + S213C + A218S + S231N | 0.3097 | **ns** |
| | **L159F** + R78K + Q309R | $1.643 \times 10^{-4}$ | *** |
| **L159F** + S213C + | **L159F** + T64A + R78K + Q309R | $1.643 \times 10^{-4}$ | *** |
| A218S + S231N | **L159F** + S213C + A218S | 0.5 | **ns** |
| | **L159F** + T64A + S213C + A218S + S231N | 0.2317 | **ns** |

[a] The statistical significance of the differences is given as follows: ns=not significant; * $p<0.05$; ** $p<0.01$; *** $p<0.001$.

**Table S5. Statistical analysis of the repertoire of substitutions with PAM250<0[a] in NS3-, NS5A-, NS5B-coding regions according to the frequency of patients in which these amino acid substitutions occur.**

| NS3 | | | |
|---|---|---|---|
| **Category 1[b]** | **Category 2** | **p-value** | **Significance[c]** |
| **10-14.9%** | > 20 | na | na |
| | 15-19.9 | 0.3141 | ns |
| | 5-9.9 | 0.2272 | ns |
| | 1-4.9 | 0.05151 | ns |
| | 0.5-0.9 | 0.4796 | ns |
| **5-9.9** | > 20 | na | na |
| | 15-19.9 | 0.5 | ns |
| | 10-14.9 | 0.7728 | ns |
| | 1-4.9 | 0.5 | ns |
| | 0.5-0.9 | 0.8416 | ns |
| **1-4.9** | > 20 | na | na |
| | 15-19.9 | 0.5 | ns |
| | 10-14.9 | 0.9485 | ns |
| | 5-9.9 | 0.5 | ns |
| | 0.5-0.9 | 0.9996 | ns |
| **0.5-0.9** | > 20 | na | na |
| | 15-19.9 | 0.3021 | ns |
| | 10-14.9 | 0.5204 | ns |
| | 5-9.9 | 0.1584 | ns |
| | 1-4.9 | 0.0003883 | *** |
| **NS5A** | | | |
| **Category 1[b]** | **Category 2** | **p-value** | **Significance[c]** |
| **1-4.9** | > 20 | 0.5 | ns |
| | 15-19.9 | 0.5 | ns |
| | 10-14.9 | 0.303 | ns |
| | 5-9.9 | 0.08192 | ns |
| | 0.5-0.9 | 0.9957 | ns |
| **0.5-0.9** | > 20 | 0.3207 | ns |
| | 15-19.9 | 0.3207 | ns |
| | 10-14.9 | 0.1038 | ns |
| | 5-9.9 | 0.00652 | ** |
| | 1-4.9 | 0.00427 | ** |
| **NS5B** | | | |
| **Category 1[b]** | **Category 2** | **p-value** | **Significance[c]** |
| **> 20** | 15-19.9 | 0.5 | ns |
| | 10-14.9 | 0.5 | ns |
| | 5-9.9 | 0.3007 | ns |
| | 1-4.9 | 0.5 | ns |
| | 0.5-0.9 | 0.5 | ns |

| | | | |
|---|---|---|---|
| **1-4.9** | > 20 | 0.5 | ns |
| | 15-19.9 | 0.5 | ns |
| | 10-14.9 | 0.5 | ns |
| | 5-9.9 | 0.1285 | ns |
| | 0.5-0.9 | 0.9893 | ns |
| **0.5-0.9** | > 20 | 0.5 | ns |
| | 15-19.9 | 0.5 | ns |
| | 10-14.9 | 0.406 | ns |
| | 5-9.9 | 0.03322 | * |
| | 1-4.9 | 0.01068 | * |

[a] PAM250 substitution matrix values are taken from Feng and Doolittle (Methods in Enzymology 266: 368-382, 1996): negative values low acceptability.

[b] Categories without substitutions with PAM250<0 not considered for the statistical analysis.

[c] The statistical significance of the differences is given as follows: na=not applicable; ns=not significant; * p<0.05; ** p<0.01; *** p<0.001.

**Table S6. Statistical analysis for the difference between HRS occurrence in basal samples of our cohort and in cohort of patients who achieved sustained virological response.**

| | **G1a** | | **G1b** | |
|---|---|---|---|---|
| **HRS** | **p-value** | **Significance[a]** | **p-value** | **Significance[a]** |
| **T64A** | $5.30 \times 10^{-9}$ | ***** | 0.5 | **ns** |
| **R78K** | 0.3387 | **ns** | na | **na** |
| **S213C** | - | **-** | 0.00428 | **** |
| **A218S** | na | **na** | $4.48 \times 10^{-5}$ | ***** |
| **S231N** | $9.05 \times 10^{-3}$ | **** | $8.99 \times 10^{-2}$ | **ns** |
| **Q309R** | $4.86 \times 10^{-2}$ | *** | 0.3162 | **ns** |

[a] The statistical significance of the differences is given as follows: na=not applicable; ns=not significant; * p<0.05; ** p<0.01; *** p<0.001.

### Figure S1

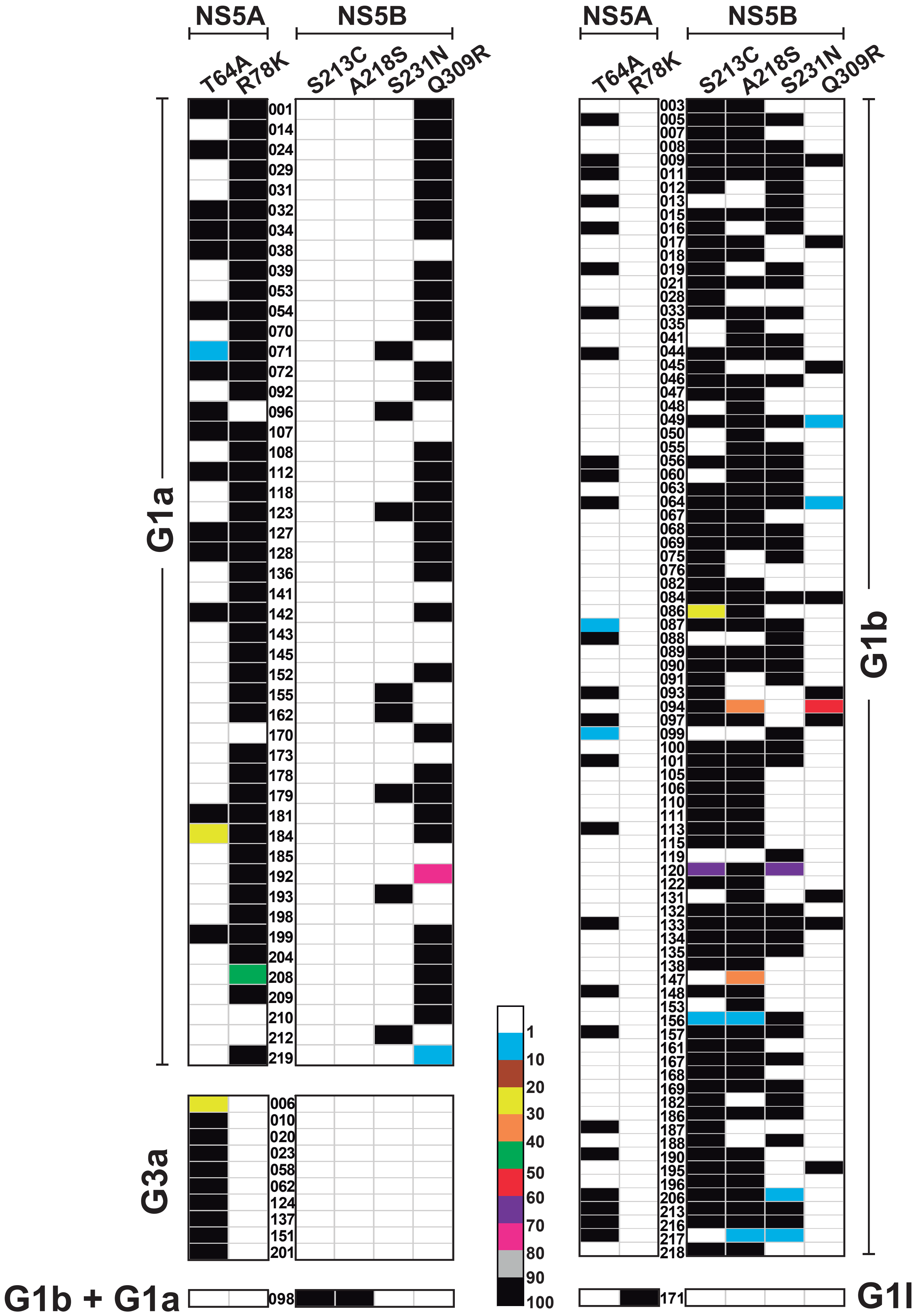
NS5A
NS5B
T64A
R78K
S213C
A218S
S231N
Q309R
G1a
G3a
G1b + G1a
G1b
G1l
1
10
20
30
40
50
60
70
80
90
100

**Figure S1. Heat map the frequency of HRS within the HCV mutant spectra of individual HCV samples at treatment failure.** The HRS and proteins NS5A, and NS5B are indicated at the top. The HCV subtype is indicated on the side of each box. Each horizontal alignment of dots represents one of the total 145 patients carrying at least one HRS (ordinate). Each vertical dot alignment corresponds to an HRS; the substitution frequency in a sample (given by the proportion of reads with the relevant mutation) is denoted by the dot color: black (90-100%), grey (80-90%), pink (70-80%), purple (60-70%), red (50-60%), green (40-50%), orange (30-40%), yellow (20-30%), brown (10-20%), blue (1-10%) and white (<1%, below the limit of detection). The ID of each patient is indicated between the boxes, and the clinical history was described in Chen et al., Antiviral Research (2020) 174: 104694.

### Figure S2

A

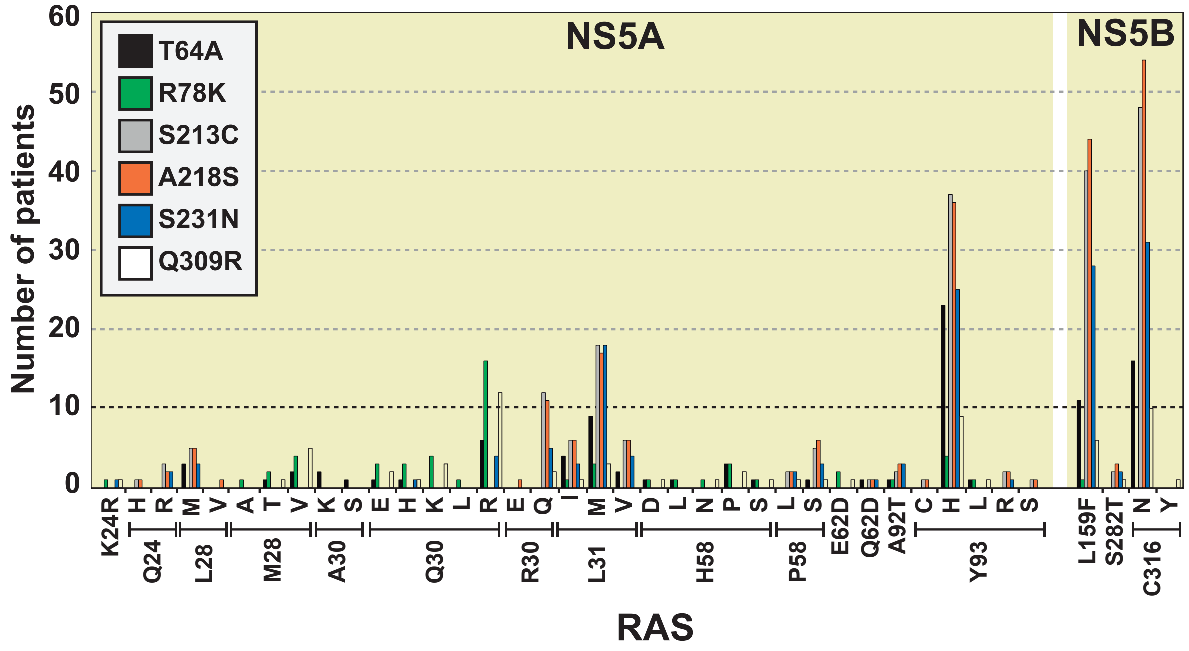


B

| HRS \ RAS | | | NS5A Q30R | NS5A R30Q | NS5A L31M | NS5A Y93H | NS5B L159F | NS5B C316N |
|---|---|---|---|---|---|---|---|---|
| | | | 19 | 13 | 26 | 74 | 46 | 54 |
| NS5A | T64A | 54 | 6 | | 9 | 23 | 11 | 16 |
| NS5A | R78K | 45 | 16 | | 3 | 4 | 1 | |
| NS5B | S213C | 72 | | 12 | 18 | 37 | 40 | 48 |
| NS5B | A218S | 70 | | 11 | 17 | 36 | 44 | 54 |
| NS5B | S231N | 57 | 4 | 5 | 18 | 25 | 28 | 31 |
| NS5B | Q309R | 47 | 12 | 2 | 3 | 9 | 6 | 10 |

C

| HRS combinations \ RAS | | | NS5A Q30R | NS5A R30Q | NS5A L31M | NS5A Y93H | NS5B L159F | NS5B C316N |
|---|---|---|---|---|---|---|---|---|
| | | | 19 | 13 | 26 | 74 | 46 | 54 |
| G1a | R78K + Q309R | 17 | 7 | | | 2 | | |
| G1a | T64A + R78K + Q309R | 13 | 3 | | 1 | 1 | | |
| G1b | S213C + A218S | 19 | | 6 | 3 | 10 | 12 | 14 |
| G1b | S213C + A218S + S231N | 18 | | 2 | 8 | 6 | 13 | 14 |
| G1b | T64A + S213C + A218S + S231N | 11 | | | 1 | 6 | 9 | 9 |

**Figure S2. Association between HRS and RASs in HCV from patients who failed therapy. (A)** Number of patients carrying each HRS and each RAS in NS5A and NS5B. **(B)** Heat map of most represented RAS (at least ten patients) associated with individual HRS. Number of patients carrying each substitution is indicated in the grey box. **(C)** Heat map of RAS (at least five patients) associated with the most represented combinations of HRS in the cohort. Number of patients carrying each RAS or HRS combination is indicated in the grey box.

#### Figure S3

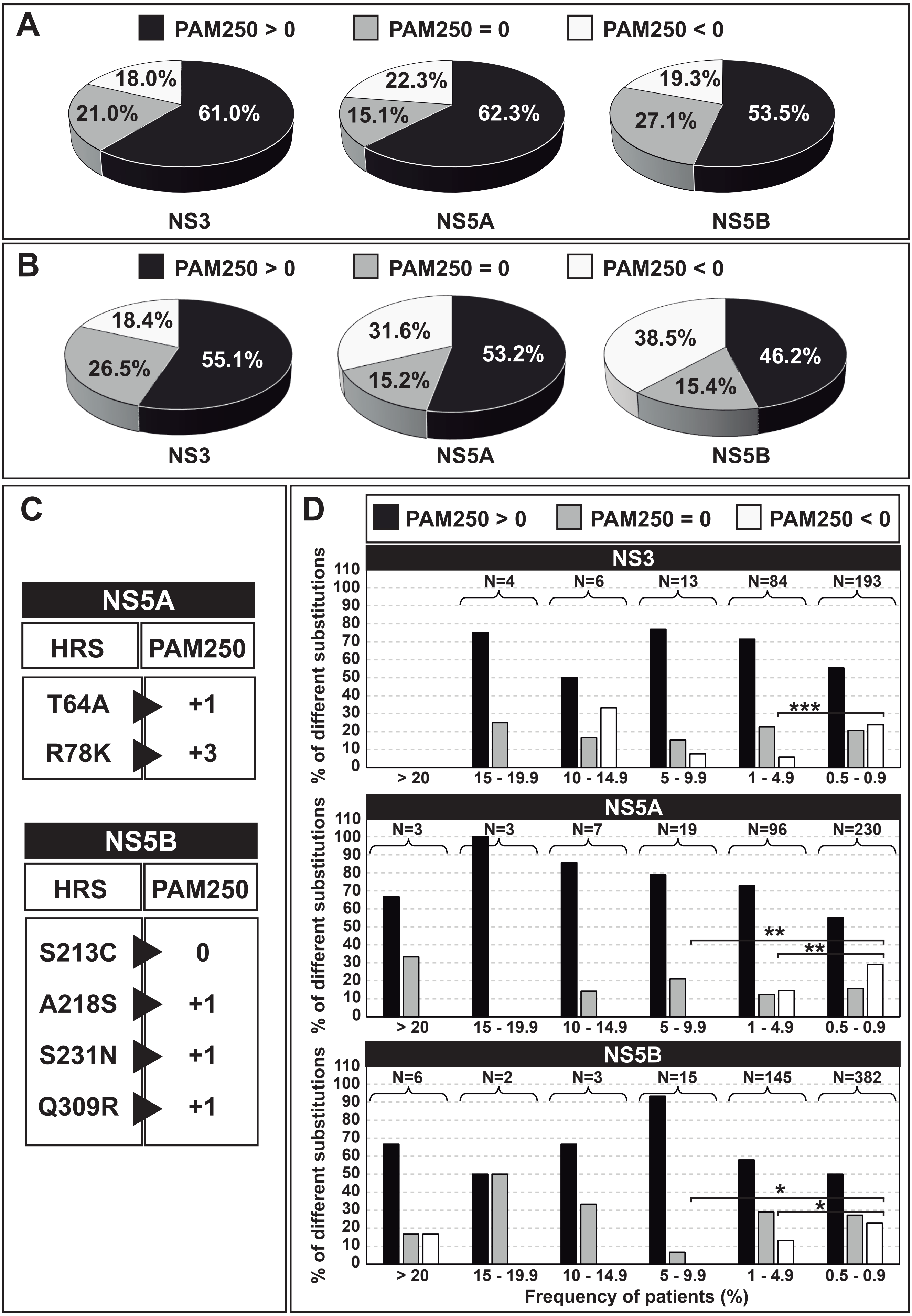

A
PAM250 > 0
PAM250 = 0
PAM250 < 0
18.0%
21.0%
61.0%
22.3%
15.1%
62.3%
19.3%
27.1%
53.5%
NS3
NS5A
NS5B
B
PAM250 > 0
PAM250 = 0
PAM250 < 0
18.4%
26.5%
55.1%
31.6%
15.2%
53.2%
38.5%
15.4%
46.2%
NS3
NS5A
NS5B
C
NS5A
HRS
PAM250
T64A
+1
R78K
+3
NS5B
HRS
PAM250
S213C
0
A218S
+1
S231N
+1
Q309R
+1
D
PAM250 > 0
PAM250 = 0
PAM250 < 0
NS3
N=4
N=6
N=13
N=84
N=193
***
NS5A
N=3
N=3
N=7
N=19
N=96
N=230
**
**
NS5B
N=6
N=2
N=3
N=15
N=145
N=382
*
*
% of different substitutions
> 20
15 - 19.9
10 - 14.9
5 - 9.9
1 - 4.9
0.5 - 0.9
Frequency of patients (%)

**Figure S3. Distribution of substitution repertoire according to the PAM250 matrix.** **(A)** Distribution of amino acid substitutions in HCV-infected patients failing DAA-based therapies. **(B-C)** Same as (A) but with RAS and HRS, respectively. **(D)** Percentage of amino acid substitutions according to PAM250 category in NS3, NS5A, NS5B according to the frequency of patients in which these amino acid substitutions occur. Patient frequency groups are indicated in abscissa, and the percentage of different amino acid substitutions in each group is given in ordinate. * $p<0.05$; ** $p<0.01$; *** $p<0.001$; proportion test.
